# Engineering growth-coupled metabolic biosensors for disease prognosis and diagnosis using full growth trajectories

**DOI:** 10.64898/2026.08.04.740108

**Authors:** Paul Ahavi, Thi-Ngoc-An Hoang, Philippe Meyer, Olivier Epaulard, Audrey Le Gouellec, Jean-Loup Faulon

## Abstract

Although metabolomics has shown considerable promise for biomarker discovery, and the development of diagnostic and prognostic applications, its translation into routine clinical practice remains limited by analytical complexity, cost, throughput, and standardization challenges. These limitations underscore the need for complementary tools, particularly in resource-limited settings. In this study, we developed a workflow for the engineering and characterization of growth-coupled metabolic sensors capable of disease detection (healthy vs. infected) and outcome prediction (mild vs. severe), which we illustrated using COVID-19 as a proof-of-concept application. We first generated a biomarker-guided library of 34 candidate sensors leveraging both auxotrophic phenotypes and less stringent metabolic dependencies. We then screened the library against patient plasma pools, identifying 19 sensor candidates with diagnostic and/or prognostic potential, including 14 with prognostic potential. Lastly, a selected subset of candidates was further evaluated on a patient cohort using two newly developed analytical frameworks designed to extract additional information from bacterial growth curves. The best-performing sensors achieved a balanced accuracy of 0.88*±* 0.06 for prognostic prediction (outer-test AUC = 0.89, 5-fold cross-validation, *n* = 37) and 1.00 for diagnostic classification (outer-test AUC = 1.00, 5-fold cross-validation, *n* = 56). Collectively, these findings establish a proof of concept for translating disease-associated plasmatic metabolic signatures into low-cost, growth-coupled biosensors with diagnostic and prognostic capabilities.

## Introduction

As metabolomics-based biomarker discovery expands across a wide range of diseases, promising diagnostic, prognostic, and precision-medicine applications are beginning to emerge [1–5]. The high cost, limited throughput, and technical complexity of current metabolomics workflows remain however major barriers to their implementation in routine clinical practice, particularly in resource-limited settings [2, 6–8]. Although efforts have been made to translate identified disease biomarkers into more accessible diagnostic solutions, such as enzymatic assays and immunoassays [2, 6], these alternatives remain highly metabolite-specific and lack the breadth of molecular coverage to represent a generalizable solution. These limitations highlight the need for adaptable, comprehensive and affordable complementary solutions, that can extend the clinical reach of metabolomics-based diagnostic tools.

In this regard, the development of such diagnostic tools has been a longstanding endeavor in synthetic biology, notably through the engineering of diverse classes of biosensors. Indeed, modern synthetic biology biosensing strategies make extensive use of bottom-up biocomputing tools, such as transcription-factor-based, protein-based and RNA-based whole-cell genetic circuits [9–11]. These classes of biosensors present several valuable characteristics: (i) they are tunable [12], (ii) able to detect a wide spectrum of potential targets [13, 14], and (iii) able to perform complex programmable operations on multiple input signals [11, 15]. Their adaptation to cell-free systems unlocked additional beneficial features, such as faster response times, improved sensitivity and deployability, and the ability to detect compounds that are toxic to microbial chassis [16].

Conversely, top-down approaches, such as growth-coupled metabolic biosensors, have received less attention owing to their limited predictability, and their challenging engineering and fine-tuning [17, 18]. However, recent advances in genome engineering [19, 20] and metabolic modeling [21, 22] have broadened, to some extent, the scope of growth-coupled designs and rendered the construction of such biosensors more accessible. More importantly, they present appealing features for clinical biosensing applications. While whole-cell circuit-based biosensors may exhibit evolutionary instability [23], or limited performances (robustness, specificity or noise) in complex media [24–26] (e.g. clinical samples), growth-coupled metabolic sensors are less prone to these issues as they rely on more stable genetic perturbations yielding consistent metabolic phenotypes. Moreover, they are particularly well suited for the detection of small metabolites derived from the primary metabolism [27], a chemical space only partially covered by the other classes of biosensors [28, 29].

The aim of this study is to demonstrate that growth-coupled metabolic biosensors can serve as low-cost, multiplexed platforms for translating metabolomics-derived biomarkers into accessible diagnostic tools in selected clinical applications. To this end, we propose a workflow for the engineering and characterization of growth-coupled metabolic sensors capable of disease detection (diagnosis) and outcome prediction (prognosis). We illustrate the utility of this strategy through the development of COVID-19 diagnostic and prognostic biosensors.

COVID-19 indeed represented a well-suited clinical case study as it caused a pandemic that highlighted the need for rapid, deployable, efficient and multiplexed diagnostic and prognostic tools [30, 31], and because several biomarkers associated with COVID-19 infection and disease outcome have already been identified through metabolomics studies [32–37]. More strikingly, we already established the ability of a bacterial strain to discriminate between COVID-19 patient groups in previous work [38]. Using a selected subset of time points from the growth curves of *E. coli* K-12 MG1655 on plasma samples from a single cohort, we successfully classified: (i) plasma samples from COVID-19-negative subjects and infected patients and (ii) pre-onset samples from patients who later developed mild or severe disease.

In the present study, we employed classification frameworks that leverage the temporal structure of bacterial growth curves rather than selected individual time points as classification features. This approach, by extracting biologically grounded growth features that reflect underlying metabolic processes, provides a more biologically interpretable representation of bacterial growth responses, substantially improving the robustness of patient classification and its potential applicability as a deployable clinical tool. Finally, our workflow establishes a rational design framework in which bacterial biosensors can be engineered so that their growth responses are tailored to the metabolic fingerprints of a target disease, providing a foundation for extending this strategy beyond COVID-19.

## Results

### Model strain *Escherichia coli* K-12 MG1655 displays strong COVID-19 diagnostic but negligible prognostic capabilities

Before implementing our workflow, we first evaluated whether metabolic engineering was required to achieve high-performance COVID-19 disease detection and outcome prediction using biosensors. To this end, we investigated the diagnostic and prognostic capabilities of the model strain *E. coli* K-12 MG1655 (Figure 1A, B, C) on our validation cohort (18 pre-onset plasma samples from mild patients, 19 pre-onset plasma samples from severe patients and 19 plasma samples from COVID-19 negative subjects).

**Figure 1:**
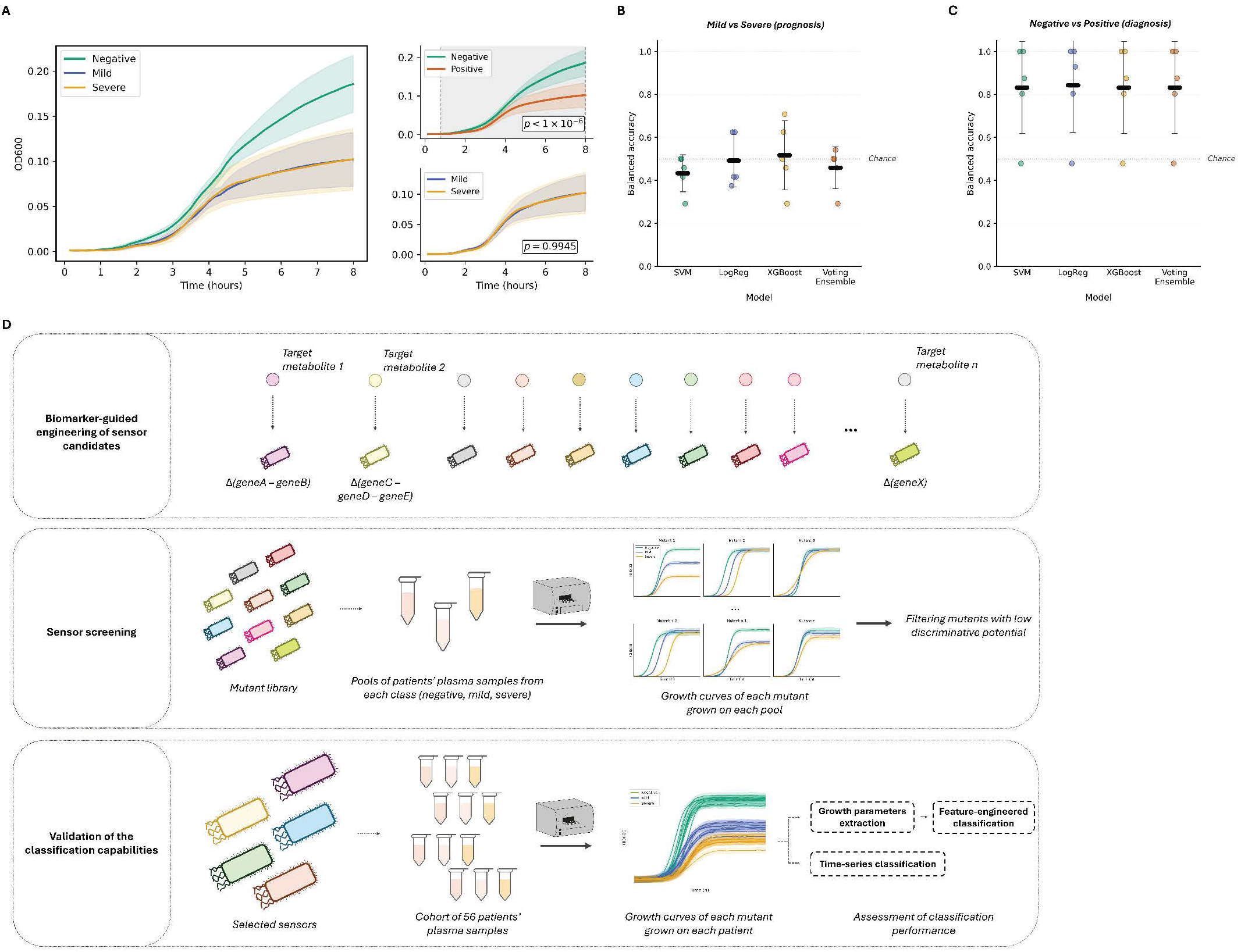
Baseline performance and study design. Panels A to C display the prognostic and diagnostic performances of *E. coli* K-12 MG1655. Panel A illustrates the mean growth curves for mild (*n* = 18, blue), severe (*n* = 19, orange), and negative (*n* = 19, green) patient samples with the standard deviation shown as shaded areas. GAMM-fitted mild and severe growth curves are shown on the upper right section, whereas GAMM-fitted negative and positive growth curves are shown on the lower right section. Benjamini-Hochberg [42]-corrected *p*-values are indicated in each section. Shaded areas represent standard deviations, and hatched areas indicate 95% confidence intervals (not readily visible in most plots owing to their narrow range). Grey windows in the GAMM representations indicate time intervals showing significant differences between conditions. Panels B and C depict the balanced accuracy for the Mild vs Severe (prognostic) and the Negative vs Positive (diagnostic) problems, respectively, using 4 classifiers: a support vector machine (SVM), logistic regression (LogReg), gradient-boosted trees (XGBoost) and an ensemble model of all 3 aforementioned classifiers. The input feature set was the maximal growth rate (*µmax*) and *OD*600*max*, as defined in the Growth parameter extraction subsection. Each dot represents one outer cross-validation fold (*n* = 5). The mean is indicated by a black bar and error bars denote the standard deviation. The grey line at a balanced accuracy of 0.5 indicates chance-level performance. All models were evaluated on identical folds. Panel D presents the study design: a library of metabolic mutants is screened to identify high-potential strains for COVID-19 prognosis and diagnosis, from which five mutants are selected and further evaluated for prognostic and diagnostic performance.

The maximal growth rate, *µ_max_*, and the *OD*600 value (biomass) at growth saturation, *OD*600*_max_*, of the model strain were determined in each patient sample, as is conventionally done in growth-coupled design analysis [18, 39, 40] (Supplementary Data 2). Indeed, the analysis of these two parameters provides improved reproducibility as they both reflect underlying biological processes and integrate information from multiple time points.

Classification performance for the prognostic (mild vs. severe patients) and diagnostic (healthy individuals vs. infected patients) tasks was evaluated using four models trained on patient growth parameters: support vector machine (SVM), logistic regression (LogReg), extreme gradient boosting algorithm (XGBoost) and an ensemble model of all three aforementioned classifiers (VotingEnsemble). The four models yielded balanced accuracies ranging from 0.83 ± 0.19 to 0.84 ± 0.2 (5-fold cross-validation) for the diagnostic task (Figure 1C) and from 0.43 ± 0.08 to 0.52 ± 0.14 (5-fold cross-validation) for the prognostic task (Figure 1B), validating our hypothesis that *E. coli* K-12 MG1655 exhibits strong diagnostic performance but negligible prognostic performance as a growth-coupled metabolic sensor within the conventional growth-coupled sensor analysis framework.

Using Generalized Additive Mixed Models [41] (GAMMs), we demonstrated that the fitted mean curves of the negative and positive classes are significantly different (*p*-value < 1 × 10*^−^*^6^, Figure 1A), whereas no significant difference was observed between the fitted mean curves of the mild and severe groups (*p*-value = 0.99, Figure 1A). More specifically, the time period significantly differing between the negative and positive groups include the exponential and stationary phases, supporting the fact that the 2 aforementioned growth parameters may enable effective diagnostic classification.

These observations corroborate the classification results, demonstrating strong COVID-19 diagnostic but negligible prognostic capabilities for *E. coli* K-12 MG1655, thereby highlighting substantial opportunities for improvement. We therefore implemented our growth-coupled sensor engineering workflow with the primary objective of isolating strains capable of prognostic classification (mild vs severe patients). A secondary objective was to enhance diagnostic performance (healthy individuals vs infected patients), as the model strain exhibited substantially lower performance than mass spectrometry-based classifiers (Table 1).

**Table 1:** Comparison of prognostic and diagnostic classification performance (AUC) between growth-coupled sensor-based classifiers and metabolomics-based models.

| Study | $n_{\text{Neg.}}$ | $n_{\text{Mild}}$ | $n_{\text{Sev.}}$ | Prognostic AUC* | Diagnostic AUC* | Independent cohort validation |
| --- | --- | --- | --- | --- | --- | --- |
| **Current work** | **19** | **18** | **19** | **0.89** | **1.00** | **N/A** |
| López-Hernández et al. [36] | 39 | 42 | 40 | 0.764 | 0.947** | N/A |
| Shen et al. [32] | N/A | 18 | 13 | 0.957*** | N/A | Correct: 7/10 |
| Roberts et al. [34] | N/A | 71 | 49 | 0.84 | N/A | AUC = 0.76 |
| Correia et al. [35] | 57 | 22 | 10 | 0.799 | 0.924 | N/A |
| Sindelar et al. [37] | N/A | 80 | 83 | 0.88 | N/A | AUC = 0.72 |

AUC was used for comparison as it was the only performance metric reported across all considered studies. $n$ represents the number of patient samples in each class ("Neg." for negative, mild, and "Sev" for severe). Prognostic AUC and Diagnostic AUC denote the AUC achieved following cross-validation for prognostic and diagnostic prediction, respectively. Independent test set indicates whether the reported performance was additionally validated on an independent test set. \* AUC resulting from a cross-validation evaluation. \*\* Negative subjects versus positive and unhospitalized patients only. \*\*\* Signs of optimistic training performance or potential overfitting, as performance on the independent test set is substantially lower. The input feature set also includes proteomics data.

### Overview of the growth-coupled metabolic biosensor engineering workflow applied to COVID-19

In order to engineer biosensors with improved capabilities for COVID-19 detection and outcome prediction, we designed a library of 34 growth-coupled metabolic candidate sensors based on the diagnostic and prognostic biomarkers identified in previous metabolomics studies (Supplementary File 1, Figure 1D). We then selected the more readily targetable biomarkers, namely amino acids, nucleobases, vitamins, polyamines and related derivatives.

Each sensor candidate was designed to target one metabolite or metabolite class. Growth medium engineering was also conducted in a personalized manner: for each sensor candidate, plasma samples were supplemented with amino acid and nucleobase mixtures to sustain steady growth and ensure optimal class separation without modifying the concentration of the corresponding target metabolites (cf. Growth medium engineering subsection). The design of the candidate sensors relied not only on auxotrophic phenotypes, but also on less stringent metabolic dependencies to demonstrate that molecular coverage of growth-coupled biosensors can be broadened to non-essential intracellular metabolites.

Candidate sensors were subsequently screened for diagnostic and prognostic potential on a limited set of patient sample pools. The clinical capabilities of five selected candidates were then validated and further characterized on a larger patient cohort (Figure 1D).

### Screening phase retained 19 candidate sensors with either diagnostic or prognostic potential

Sensor screening consisted of culturing each candidate in three plasma sample pools of each class (mild, negative, severe) supplemented with the corresponding nutrient mix (Figure 1D). Diagnostic and prognostic potential was evaluated by comparing group-wise mean trajectories for the negative versus positive classes and the mild versus severe classes, respectively, using Generalized Additive Mixed Models [41] (*α* = 0.1). This pooling strategy was implemented as screening every mutant across all patient samples in the cohort was not feasible due to sample volume limitations. Moreover, this approach enabled the reduction of the amount of plasma required for screening, preserving more patient samples for the sensor validation phase. However, it also reduced the statistical resolution of the screen, allowing only the identification of potential sensors requiring further confirmation. In addition, focusing solely on group-wise mean trajectory differences to identify classification potential excluded candidate sensors generating individual growth patterns exploitable by machine-learning classifiers.

Out of the 34 candidates tested, 13 were screened out due to lack of growth on plasma, residual growth, or uninterpretable growth profiles (Supplementary Figure 1). The GAMM analysis identified 19 candidate sensors with classification potential: (i) five with both prognostic and diagnostic potential (Figure 2A), nine with prognostic potential only (Figure 2B), and 5 with diagnostic potential only (Supplementary Figure 2A). Among them, six also showed potential for multiclass classification between all three classes (Supplementary Figure 2B). Two candidates failed the screening, displaying neither prognostic nor diagnostic potential (Supplementary Figure 2C).

**Figure 2:**
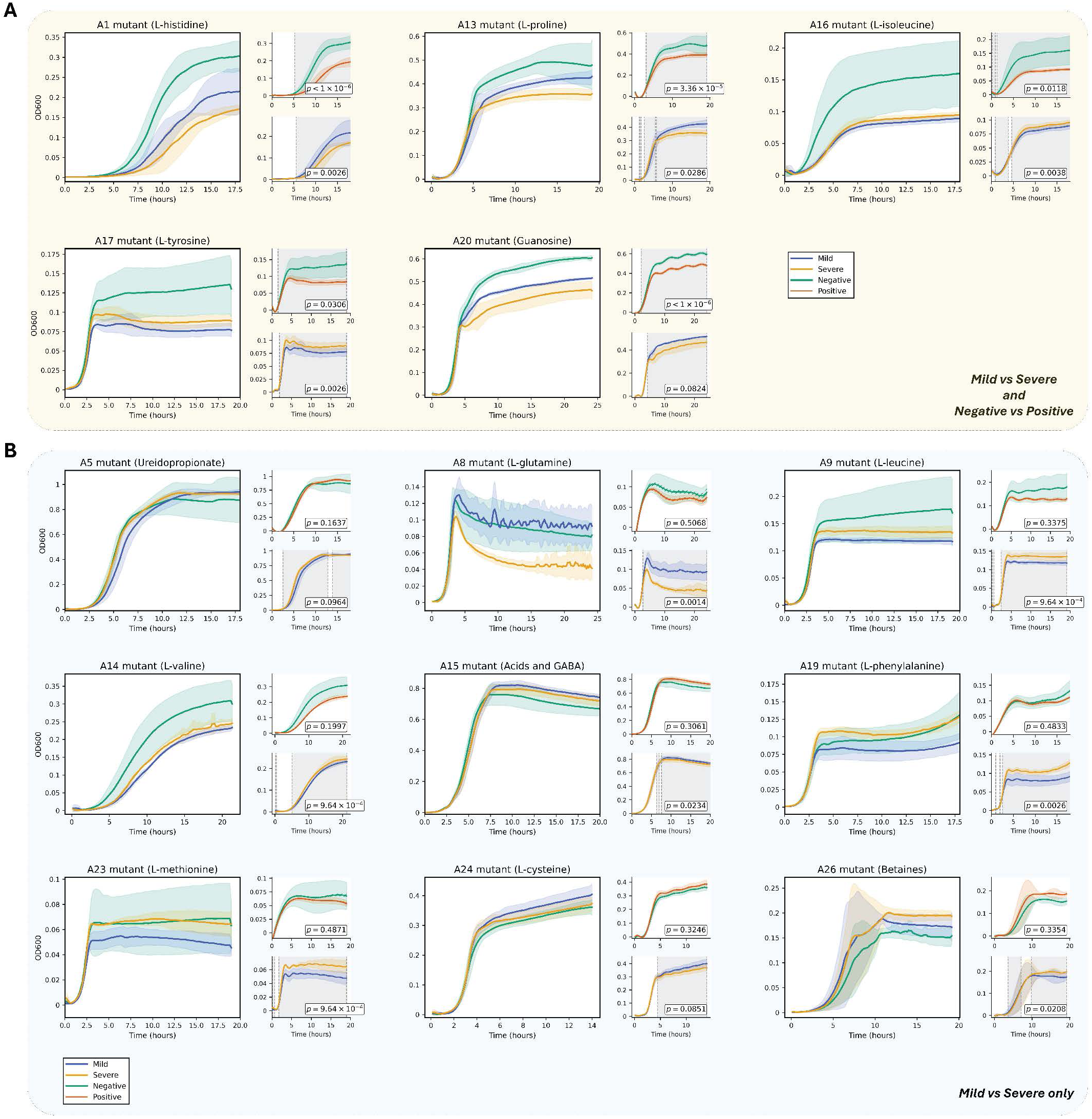
Metabolic candidate sensors with COVID-19 prognostic potential. Prognostic and diagnostic potential was assessed using generalized additive mixed models [41] (GAMMs) applied to growth curves obtained from mutants cultured in mild, severe, and negative plasma sample pools (*n* = 3, *α* = 0.1). Panel A shows mutants with both prognostic and diagnostic potential, whereas Panel B shows mutants with prognostic potential only. For each mutant, mean growth curves are indicated in the left section, GAMM-fitted mild and severe growth curves in the upper-right section and GAMM-fitted negative and positive growth curves in the lower-right section. Mild curves are colored in blue, severe curves in orange, negative curves in green and positive curves in red. Corresponding *p*-values (corrected using Benjamini-Hochberg procedure [42] to account for multiple testing) are indicated in each section. Shaded areas represent standard deviations, and hatched areas indicate 95% confidence intervals (not readily visible in most plots owing to their narrow range). Grey windows in the GAMM representations indicate time intervals showing significant differences between the two conditions.

Group-wise differences seem to occur at minimum during stationary phase for all screen-positive mutants, consistent with a growth phenotype associated with nutrient depletion or acidification (in the case of A15). In most cases, however, the observed divergence period is extended partially or throughout the exponential phase (Figure 2, Supplementary Figure 2). The involvement of several growth phases could potentially enhance classification robustness as it could provide more opportunities to detect discriminative phenotypes.

### Validation stage confirmed the prognostic and diagnostic capabilities of five selected mutants

We further confirmed the prognostic and diagnostic capabilities of 5 selected candidate sensors (A1, A5, A15, A19, A28) on a larger cohort, thereby increasing the statistical power of the analysis (Figure 3A). To that effect, the five selected mutants were cultured in individual plasma samples from all remaining patients of the cohort (*n_Mild_*= 18, *n_Severe_*= 19, *n_Negative_*= 19; Supplementary Figures 13 to 18). It is noteworthy that the patient samples used to prepare the plasma pools during screening were excluded from the validation cohort and that this is the same validation cohort that was used to previously assess the classification capabilities of the model strain *E. coli* K-12 MG1655 (Figure 1).

**Figure 3:**
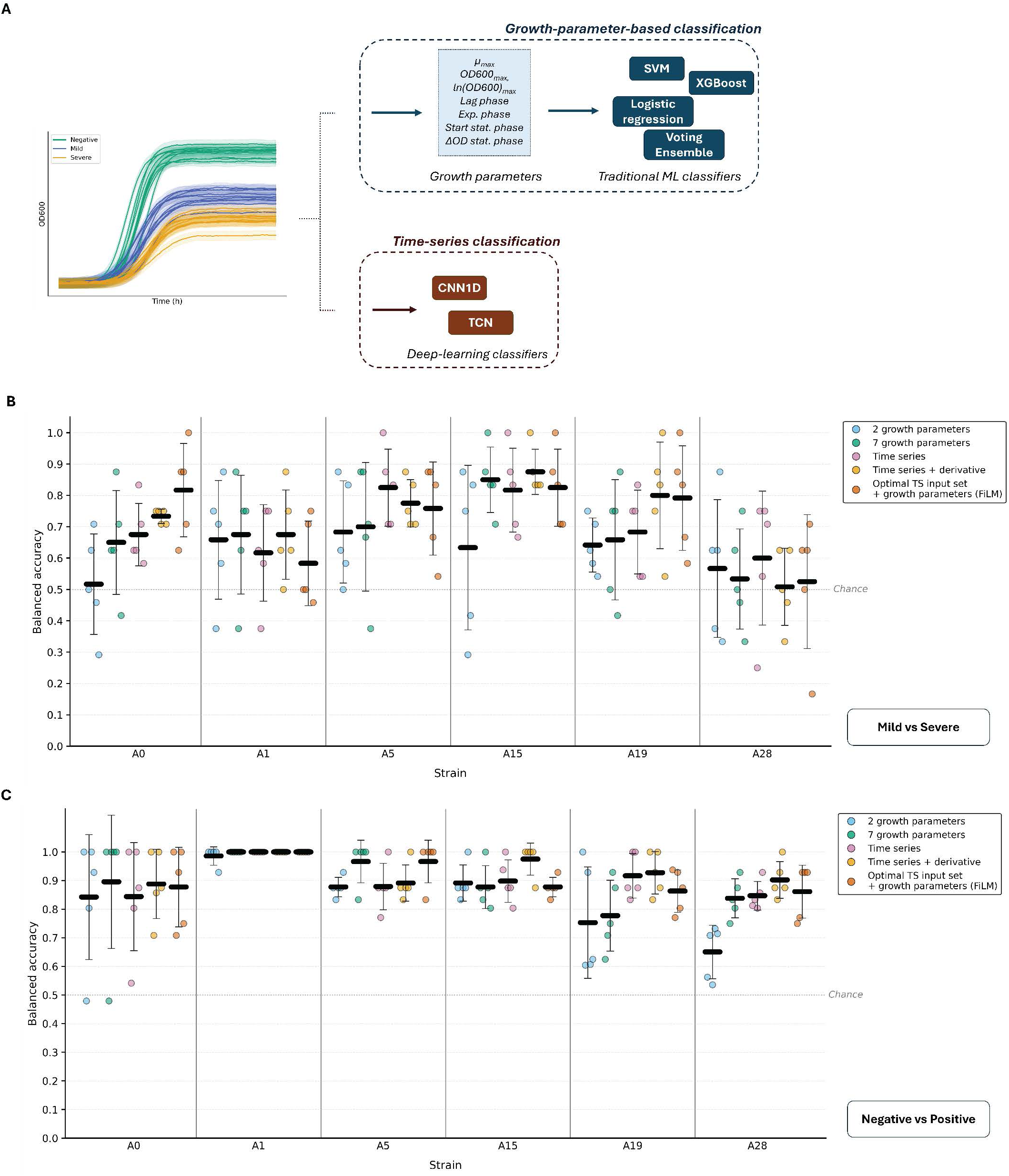
Prognostic and diagnostic classification using the five selected metabolic sensor candidates. Panel A presents the two approaches compared in this study: (i) conventional growth-parameter-based classification using machine learning classifiers, including a support vector machine (SVM), logistic regression (LogReg), gradient-boosted trees (XGBoost), and an ensemble model and (ii) time-series classification using deep-learning classifiers, including a one-dimensional convolutional neural network (CNN1D) and a temporal convolutional network (TCN). Panels B and C show, for *E. coli* K-12 MG1655 and the five selected metabolic candidate sensors, the highest balanced accuracy obtained for prognosis (mild vs severe classification) and diagnosis (negative vs positive classification), respectively, using different feature sets: (i) 2 growth parameters, (ii) 7 growth parameters, (iii) time series, (iv) time series combined with the first derivative, and (v) time series combined with growth-parameter conditioning through feature-wise linear modulation [43] (FiLM). Each dot represents one outer cross-validation fold (*n* = 5). The mean is indicated by a black bar and error bars denote the standard deviation. The grey line at a balanced accuracy of 0.5 indicates chance-level performance.

We trained four machine learning classifiers, namely a support vector machine (SVM), a logistic regression (LogReg), gradient-boosted trees (XGBoost), and an ensemble model, on the two growth parameters (*µ_max_* and *OD*600*_max_*) extracted from the mutant growth curves (Figure 3A).

The results corroborated the screening findings regarding the outcome prediction task (Figure 3B), as classifiers based on A1, A5, A15, and A19 displayed moderate prognostic performance (between 0.63 ± 0.23 and 0.68 ± 0.15), while A28-based models exhibited negligible prognostic capabilities. However, the screening appeared overly stringent for diagnostic (Figure 3C) and multiclass potential (Supplementary Figure 4), failing to identify the strong diagnostic performance of A5, A15 (up to 0.89 ± 0.05, comparable to the model strain), the moderate performance of A19, and the moderate multiclass classification capabilities of A5 and A19. This is likely attributable to the limited screening sample size (*n* = 3 per class). Of note, A1-based classifiers reached a balanced accuracy of 0.99 ± 0.03, corresponding to a single misclassification (Supplementary Figure 6).

Although our workflow successfully identified prognostic biosensors, their performance remained insufficient to meet clinical requirements or match that of other mass spectrometry-based classifiers (Table 1). Growth-coupled metabolic sensors often exhibit non-canonical growth curves. To better capture these growth dynamics, we explored more comprehensive analytical frameworks that leverage information beyond that captured by analyses based solely on *µ_max_* and *OD*600*_max_*.

### An extended seven-growth-parameter feature set enables strain-dependent improvements in classification performance

Our first approach consisted of extending the conventional two-parameter growth analysis by extracting five additional biologically relevant features from growth curves. The five additional parameters are: (i) maximal ln-transformed OD600, *ln*(*OD*600)*_max_*; (ii) lag phase duration, (iii) exponential phase duration; (iv) onset time of the stationary phase and (v) difference between maximal and final OD600, *OD*600*_max_* − *OD*600*_final_* (cf. Growth parameter extraction subsection). All five parameters are biologically grounded and capture distinct aspects of bacterial growth dynamics. Lag phase duration reflects the time required for a strain to adapt to the culture medium before entering exponential growth. The maximum ln-transformed OD600, *ln*(*OD*600)*_max_*, and the duration of the exponential phase provide information about the deceleration phase, when such a phase is observed. This transitional stage precedes the stationary phase and is characterized by a progressive reduction in the specific growth rate. The onset time of the stationary phase corresponds to the time required for bacterial growth to alter the medium to the point where growth becomes limiting, for instance through nutrient depletion, acidification, or accumulation of metabolic by-products. Finally, the difference between maximum and final OD600, *OD*600*_max_* − *OD*600*_final_*, reflects population dynamics during the stationary phase (biomass stability, residual biomass accumulation, or cell death).

Using the extended seven-parameter feature set improved both the prognostic and diagnostic performance of *E. coli* K-12 MG1655 (A0), with balanced accuracies reaching 0.65 ± 0.15 and 0.90 ± 0.21 (5-fold cross-validation), respectively. Moreover, when evaluated on identical cross-validation folds, classifiers trained on the extended feature set achieved higher or comparable balanced accuracies across all six strains and classification tasks (Figure 3B, C, Supplementary Figure 4). Improvements were observed for A0 and A15 in prognostic prediction (balanced accuracy increases of 0.13 and 0.22, respectively), for A5 and A28 in diagnostic classification (increases of 0.09 and 0.19, respectively), and for A0, A15, and A28 in multiclass classification (increases of 0.13, 0.13, and 0.19, respectively). Performance remained comparable between the two feature sets for the other strains. These results suggest that the seven-parameter analysis more effectively captures strain-specific discriminative information across diverse growth profiles and genetic backgrounds.

Under this extended analytical framework, A15 emerged as the best-performing prognostic biosensor, achieving a balanced accuracy of 0.85 ± 0.09 (5-fold cross-validation), whereas the other mutants exhibited prognostic performance comparable to that of A0. Regarding the diagnostic task, A1-based models correctly classified all 56 individuals in the cohort (Figure 3, Supplementary Figure 6), while A5-based models reached a balanced accuracy of 0.96 ± 0.07 with a single misclassification (5-fold cross-validation).

### Full growth curve analysis generally yields superior classification performance compared with growth-parameter-based approaches

In order to investigate whether the prognostic performance of the strain classifiers could be further improved by leveraging the full information contained in the growth curves, we explored a second approach based on time-series classification. The full growth curves were used as input to two deep-learning models (Figure 3A): a one-dimensional convolutional network (CNN1D) and a temporal convolutional network (TCN). Indeed, analysis of the full time series may reveal additional temporal patterns or correlations that are not captured by growth parameters. We tested two different input feature sets: (i) full time series (TS) and (ii) full time series combined with its first derivative (TS+), which might provide additional information about local behaviors.

Time-series frameworks improved prognostic and multiclass prediction performance for four strains (out of 6 total), with balanced accuracy increases of up to 0.14 and 0.16, respectively (Figure 3B, Supplementary Figure 4). Of note, classifiers trained on TS+ input sets demonstrated superior or comparable performance relative to growth-parameter-based classification across tested sensors for the prognostic and multiclass tasks. No universal optimal input set emerged between TS and TS+; the best-performing configuration was strongly strain- and task-dependent.

Results were more mixed regarding diagnostic classification, with performance improvements observed for three sensors but performance reduction for another (Figure 3C). This outcome is not surprising, as growth-parameter classifiers already capture sufficient information to achieve strong diagnostic accuracies. In that case, time-series analysis might introduce noise or non-informative features.

Motivated by the partial complementarities between the patient-level predictions of growth-parameter and time-series classifiers (Supplementary Figures 5 to 7), we finally tested a mixed approach between time-series and growth parameters. Multimodal classifiers are trained on the optimal time-series input set (TS or TS+) together with growth parameters as contextual inputs using Feature-wise Linear Modulation [43] (FiLM). In this framework, growth parameters modulate feature extraction from the time-series data. With the exception of two cases (prognosis using A0 and diagnosis using A5), this strategy consistently underperformed across all strains and classification tasks, probably owing to noise or information redundancy between the features extracted from the time series and the growth parameters.

Overall, we identified growth-coupled metabolic sensors capable of strong prognostic and perfect diagnostic classification across the cohort, achieving balanced accuracies of 0.88 ± 0.06 and 1.00 (5-fold cross-validation), respectively. Among the five tested mutants, one (A15) outperformed the model strain A0 for prognostic prediction, while three achieved superior diagnostic performance, with perfect classification for A1 and near-perfect classification for A5 and A15 (one misclassification each). Notably, two strains (A5 and A15) combined both strong prognostic and near-perfect diagnostic capabilities.

### Multistrain models do not improve prognostic performance

Building on partial complementarities between patient-level predictions of strains best-performing models (Supplementary Figures 5 to 7), we investigated whether multistrain models, combining input sets from several strains, could further improve prognostic performance. Two fusion configurations were evaluated: (i) early-fusion classifiers (CNN1D and TCN), trained on the optimal input sets of each strain, and (ii) a late-fusion classifier consisting of a soft-weighted ensemble of the best-performing base models from each participating strain (Supplementary Figure 8).

No multistrain combination outperformed the best-performing single-mutant model (A15). Early-fusion classifiers consistently underperformed relative to late-fusion models. Within late-fusion configurations, models reaching performance levels similar to the standalone A15 classifier are the A0 + A1 + A5 model and all classifiers incorporating the A15 base model. Examination of ensemble weights revealed that the A5 base model contributed most to the A0 + A1 + A5 ensemble, whereas the A15 base model dominated all ensembles incorporating A15 (Supplementary Data 3).

Analysis of patient-level predictions showed that correct predictions arising from complementary information between strains were counterbalanced by misclassifications stemming from conflicting predictions across strains (Supplementary Figures 9 to 11). Overall, the noise introduced by multistrain fusion models appeared to outweigh the associated gain in discriminative information, possibly owing to the small cohort size.

### Group-wise mean trajectory differences support the biological relevance of the patterns underlying classification

Finally, statistical analyses were conducted on the full growth curves (GAMMs [41]) and on the seven growth parameters (Welch’s *t*-test [44]) of the validation cohort in order to assess group-wise differences in mean dynamics and therefore investigate the existence of group-level discriminative signals that would support the classification results.

Both strategies identified significant differences in the mean trajectories between the negative and positive groups and between the mild, severe and negative classes for all six sensor strains (Figure 4, Supplementary Figure 12). Only A1, A5 and A15 mutants showed significant differences in mean profile between the mild and severe groups (Figure 4). These findings are largely consistent with the classification results and suggest that the patterns captured by strain classifiers are biologically meaningful, further reinforcing the robustness of the approach. However, no significant mean dynamics differences were identified between the mild and severe groups for A0 and A19, with GAMM *p*-values of 0.99 and 0.91, respectively. The fact that A0- and A19-based models still achieved a strong prognostic classification performance (0.82 ± 0.13 and 0.8 ± 0.15 in a 5-fold cross-validation evaluation, respectively), despite the absence of significant differences between the two classes, suggests that information underlying prognostic classification may be encoded in individual growth dynamics rather than in group-level mean behaviors for these two cases.

**Figure 4:**
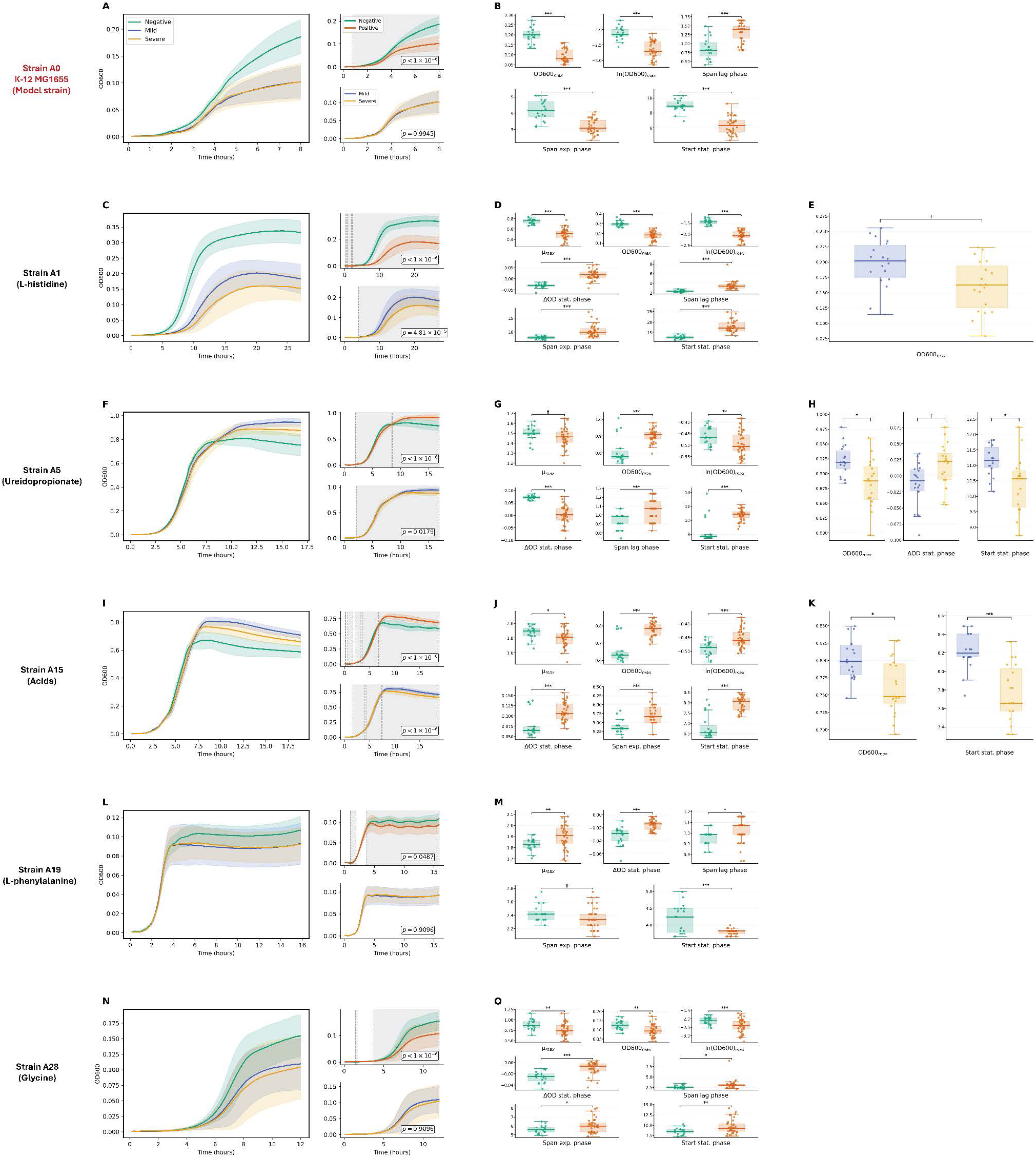
Confirmation of group-wise mean trajectory differences between mild versus severe and negative versus positive classes for the five selected metabolic sensors. For each metabolic sensor, mean growth curves for the mild (*n* = 18), severe (*n* = 19) and negative (*n* = 19) patient samples are illustrated in the left section of the first column of panels (A, C, F, I, L, N). GAMM-fitted mild and severe growth curves are shown on the upper right section for each metabolic sensor (A0, A1, A5, A15, A19 and A28 strains) and GAMM-fitted negative and positive growth curves on the lower right section (*α* = 0.1). Mild curves are colored in blue, severe curves in orange, negative curves in green and positive curves in red. Corresponding *p*-values (corrected using Benjamini-Hochberg procedure [42] to account for multiple testing) are indicated in each section. Shaded areas represent standard deviations, and hatched areas indicate 95% confidence intervals (not readily visible in most plots owing to their narrow range). Grey windows in the GAMM representations indicate time intervals showing significant differences between conditions. The second column of panels (B, D, G, J, M, O) shows boxplots of the growth parameters that differed significantly between the negative and positive groups for each mutant according to a Welch’s *t*-test [44] (*α* = 0.1). Each dot represents a growth curve acquired on an individual patient sample. *p*-values (corrected using Benjamini-Hochberg procedure [42] to account for multiple testing) are indicated as follows: *p <* 0.001 (***), *p <* 0.01 (**), *p <* 0.05 (*) and *p <* 0.1 (*†*). The third column of panel (H, K) shows boxplots of the growth parameters that differed significantly between the mild and severe groups according to a Welch’s *t*-test [44] (*α* = 0.1). No boxplots are shown for strains A0, A19, and A28 because no significantly different growth parameters were identified between the mild and severe groups for these strains.

More specifically, discriminative growth phases between classes identified by the GAMM analysis are corroborated by the corresponding growth parameters, although the GAMM analysis occasionally identified additional differences (e.g. between the mild and severe classes for A1; Figure 4). These results might explain why time-series frameworks generally outperformed growth-parameter-based classification.

Significant differences in some of the 5 newly introduced growth parameters were observed across all classification tasks, further demonstrating that these novel parameters contain discriminative information. These observations also confirmed that the seven-parameter framework generally captures more informative signal than the conventional two-parameter-based classification across all six genetic contexts tested, explaining the improved classification results.

## Discussion

As part of a broader effort to develop new low-cost, multiplexed complementary solutions to metabolomics-driven diagnostic and prognostic tools, we designed a workflow for the engineering and the characterization of growth-coupled metabolic biosensors capable of disease detection and outcome prediction. We successfully applied it to the COVID-19 case study, with biosensor systems achieving a balanced accuracy of 0.88 ± 0.06 (outer-test AUC = 0.89, 5-fold cross-validation, *n* = 37) for prognostic prediction and 1.00 (outer-test AUC = 1.00, 5-fold cross-validation, *n* = 56) for diagnostic classification. These scores are comparable to those reported in other mass spectrometry-based studies (Table 1), supporting the translational potential of growth-coupled biosensing as a complementary prognostic and diagnostic approach in selected clinical applications. It is noteworthy, however, that RT-PCR remains the reference standard for COVID-19 diagnosis [45], and that the present study is not intended to challenge this method. Moreover, the diagnostic task evaluated in this study was restricted to the classification of healthy individuals and COVID-19 patients; future studies involving patients with non-COVID diseases will be required to assess disease specificity. More importantly, validation of this proof of concept on an independent patient cohort will be essential to further evaluate the robustness and generalizability of the proposed approach across different patient populations. That said, only five mutants were fully characterized in this study; therefore, an exhaustive performance evaluation of the remaining 14 candidate sensors exhibiting COVID-19 clinical stratification potential may reveal biosensors with superior prognostic or diagnostic performance.

The first stage of the proposed workflow consists of biomarker-guided sensor engineering, leading, in this case study, to the construction of a library of 34 growth-coupled metabolic biosensors. We demonstrated that growth-coupled sensors are not restricted to auxotrophic designs, and can also leverage more complex metabolic dependencies via the engineering of selected combinations of genetic and environmental contexts. These findings broaden the spectrum of chemical species covered by growth-coupled sensors to various non-essential metabolites, such as ureidopropionate (A5), acids (A15), or glycine betaine (A26). Notably, the A5 and A15 mutants proved to be the best-performing biosensors identified in this study when considering combined prognostic and diagnostic performance. In addition, the A26 and A30 mutants successfully passed the functional screening, demonstrating prognostic, and diagnostic classification potential, respectively.

As the number of metabolomics studies investigating disease biomarkers in human biofluids continue to progress rapidly [46, 47], this approach could be generalized across a wide range of clinical contexts, provided that the identified biomarkers fall within the scope of growth-coupled sensors. Indeed, compared to mass spectrometry, which enables precise molecule profiling and broad coverage across small metabolites, growth-coupled sensing faces two constraints: (i) it has a narrower coverage, and (ii) depending on the nature of the molecule and the chassis, detection resolution might be limited to the metabolite class rather than the individual molecule. For instance, while several *E. coli* growth-coupled sensors are able to detect individual amino acids, achieving a similar resolution for fatty acids is more difficult because many are processed through common metabolic pathways. As a result, some high-value disease biomarkers may be challenging to detect selectively. Metabolite coverage and resolution could potentially be expanded by employing alternative bacterial species as chassis, particularly as genome-editing tools for non-model organisms are being increasingly accessible [48]. Another strategy would be to rewire the metabolism of *E. coli* through enzyme engineering or stable network-level metabolic engineering, making possible phenotypical responses to previously untargetable metabolites. These perspectives naturally extend toward minimal genome [49] and artificial-cell engineering [50, 51], which could ultimately offer unprecedented flexibility in the design of metabolic networks.

The second step of the procedure consists of screening the newly engineered sensors on plasma sample pools, resulting, in this case study, in the identification of 19 candidate sensors with either prognostic or diagnostic potential (out of a total of 34), including 14 with prognostic potential. Lastly, the final stage comprises the performance validation and further characterization of a selected subset of promising candidate sensors on a patient cohort. Within the constraints of the available plasma volume for each patient, five candidates were selected in this case study, revealing two candidate sensors (A5 and A15) that displayed a better combined diagnostic and prognostic performance than the model strain *E. coli* K-12 MG1655, as well as a candidate (A1) that achieved a perfect diagnostic classification of all patients in the cohort in a 5-fold cross-validation setup. Strikingly, significant group-wise differences in mean trajectories were observed across a large majority of evaluated cases, suggesting that classification performances are rooted in biologically meaningful differences between classes, and thereby strengthening the robustness of the results. Taken together, these findings constitute a proof of concept for the ability of the proposed workflow to translate selected plasmatic metabolomic signatures into clinically actionable, high-performing prognostic and diagnostic biosensors. It is also noteworthy that all selected mutants as well as the model strain also achieved strong performance in multiclass classification across three clinically relevant groups (mild, severe, and negative), suggesting that it might be possible to predict more granular clinically relevant outcomes, such as mortality [34].

Finally, we introduced two more informative analytical frameworks for growth-coupled designs relying on: (i) an expanded growth-parameter-based strategy based on the extraction of seven biologically relevant growth parameters (*µ_max_*, *OD*600*_max_*, lag phase duration, exponential phase duration, *ln*(*OD*600)*_max_*, stationary phase onset and *OD*600*_max_* − *OD*600*_final_* difference) and (ii) a full growth curve approach using deep-learning models. Both strategies provide a more flexible and comprehensive representation of growth curves that may better fit non-canonical growth curves from metabolic sensors and thereby enrich classifier models with more biologically meaningful information. While time-series classifiers generally outcompeted models fed on growth parameters across all candidate sensors and classification tasks, the best input feature set (and thus the nature of the discriminative information) varied depending on both the strain and the classification task. We therefore propose a dynamic, strain-dependent and problem-dependent framework that will use the most informative input set in each case.

On a broader note, this study falls within an emerging direction in microbial computing, which seeks to exploit nonlinear intrinsic cellular [38] and collective [52] temporal dynamics to perform complex tasks in a robust and energy-efficient manner. Future efforts might involve the acquisition of additional easy-to-monitor phenotypes, as well as the engineering of metabolic sensor systems able to yield more complex behaviors. Indeed, mapping the discriminative biomarkers into a higher dimensional space might improve class separability [53]. Microbial consortia constitute a promising lead in this regard, as they are able to leverage both individual and collective dynamics to produce more complex phenotypes. Engineering microbial-consortia-based biosensors might enable the simultaneous sensing of multiple metabolites and therefore the detection of multivariate metabolomic signatures, which could improve the disease specificity of this biosensing approach. From a technology transfer perspective, further miniaturization could be highly beneficial to enhance portability and practicality, hence making devices more suitable for point-of-care testing [54, 55], especially in resource-limited settings. In that regard, the development of a hybrid device combining microfluidics chips and integrated systems constitutes a promising direction toward real-world implementations.

## Methods

### Code and data accessibility

All source codes (classification models, generalized additive mixed models, Welch’s *t*-test) as well as data generated in this study and required to reproduce the results are available in two Zenodo repositories [56, 57]. Source codes are also available in this GitHub repository: https://github.com/brsynth/mutant-covid.

### Strains and media

All strains, primers and media used in this study are described in the Supplementary File 1.

#### Genome engineering

The aim of this procedure was to engineer metabolic sensors displaying a distinctive growth phenotype when cultured in plasma samples from each class. To this end, mutants matching the diagnostic and prognostic biomarkers were engineered. Both diagnostic and prognostic biomarkers were identified by literature search. The list of target metabolites was narrowed down to metabolites considered potentially critical for *E. coli* as they are expected to yield strong phenotypic responses: (i) amino acids and derivatives, (ii) nucleobases and derivatives, (iii) vitamins and (iv) polyamines (cf. Supplementary File 1). Mutants were then engineered to yield a variety of responses ranging from nutrient-responsive phenotype to auxotrophy to each of these selected metabolites. The environmental context was also engineered to tailor the phenotypes obtained to the nature of the samples and to maximize growth phenotype differences between classes, as detailed in the Growth medium engineering subsection.

For each target metabolite or metabolite class, the corresponding gene deletion strategy was designed based on literature search and/or using KEGG [58] and EcoCyc [59] databases. All gene deletions carried out in *Escherichia coli* followed the procedure developed in Jiang *et al.* [60]. Briefly, this protocol uses CRISPR-Cas9 to generate double-strand breaks at the target genome locus, and the lambda red recombinase to introduce a desired mutation via homologous recombination.

The deletion strategy consisted of deleting almost the entire coding sequence of each target gene whenever possible, provided that it did not affect the regulation of neighboring genes too extensively. The latter criterion was evaluated using RegulonDB [61], by ensuring that as few transcription factor binding sites as possible were deleted. The few remaining amino acids at the 5*^′^*and 3*^′^* ends of the deleted region were not part of any catalytic site (checked using UniProt [62]) and were kept in frame; or if they were not, a stop-codon was introduced in order to prevent alterations in the regulation of downstream genes.

For each target gene, a specific N20 guide sequence, designed using CHOPCHOP [63], was cloned by inverse PCR into the pTargetF plasmid using the primers listed in Supplementary File 1. Donor DNA, consisting of a DNA fragment corresponding to approximately 500 bp upstream of the deleted region concatenated with a second fragment corresponding to approximately 500 bp downstream of the deleted region, was cloned by overlap-extension PCR using the primers listed in Supplementary File 1.

For each target gene, 500ng of donor DNA was co-transformed with 100ng by electroporation (1mm cuvettes, 1.8 kV) into competent *E. coli* K-12 MG1655 cells harboring a pCas plasmid, following the procedure described in Jiang *et al.* [60]. Transformants were selected and screened by colony PCR using the primers listed in Supplementary File 1. The positive clones were isolated and cured on plates following the protocol described in Jiang *et al.* [60]. Multiple-deletion mutants (up to 6) were obtained after several iterations of this process.

#### Growth medium engineering

*OD*600*_max_* values extracted from *E. coli* K-12 MG1655 growth curves on plasma samples were low (mean = 0.132 ± 0.066; compared with values up to 0.9 in rich medium; Supplementary Figure 13). Introducing additional genetic perturbations under these conditions could either mask growth phenotype differences across groups or even prevent bacterial growth. To overcome this limitation, plasma samples were supplemented with nutrients at concentrations indicated in Supplementary File 1. More specifically, all proteinogenic amino acids and nucleobases were supplemented for each mutant, except for the target metabolites and those likely to rescue the phenotypic effects of the corresponding genetic perturbations. This setup directly links the growth phenotype to the concentration of the target metabolite while maximizing growth differences across classes.

Additionally, plasmatic concentrations of the target metabolites across groups may not always fall within the dynamic range of the metabolic biosensors. To address this issue, each mutant was tested on three plasma dilutions during sensor screening (as described in the Sensor screening subsection).

### Cultures of *Escherichia coli* in plasma samples

#### Plasma sample collection

The plasma sample set used in this study comprised 35 samples from COVID-19 negative subjects, 34 pre-onset samples from patients who later developed a mild form of the disease, and 35 pre-onset samples from patients who later developed a severe form of the disease. Medical definitions of “mild” and “severe” are provided later in this subsection. Plasma samples were supplemented with ethylenediaminetetraacetic acid (EDTA) as anticoagulant.

Plasma samples from COVID-19 negative subjects were acquired from the French Blood Establishment (Etablissement français du sang, EFS), while samples from COVID-19 infected patients were extracted from the BIOMARCOVID cohort, a single-center, retrospective cohort study of patients hospitalized at the Grenoble-Alpes University Hospital (CHUGA) for COVID-19. This non-interventional, monocentric, retrospective study involving data and samples from human participants has been carried out in CHUGA according to French current regulation. The primary aim for that cohort was to develop a regression model that integrated clinical and biochemical parameters related to hospitalization and disease severity. The secondary aim was to identify differential metabolites and metabolic pathways between mild and severe COVID-19 outcomes using metabolomics analysis.

Inclusion criteria included hospitalization for suspected COVID-19 and a positive PCR test for SARS-CoV-2. Exclusion criteria included age under 18, transfer from another hospital, opposition to research, ICU admission on the first day, and blood sample collection more than 48 hours after hospitalization at CHUGA. Clinical evaluation recorded demographic data (age, sex, BMI), disease progression (ICU, length of stay), and outcome status (class: mild or severe).

Specialized teams assisted in acquiring clinical data. Patient outcomes were classified retrospectively using the WHO ten-category ordinal scale into six groups. Category 3: Patients hospitalized without oxygen supplementation; Category 4: *O*_2_ flow rate ≤ 2 L/min; Category 5: *O*_2_ flow rate *>* 2 L/min; Category 6 : admitted to ICU. The categories 3 and 4 were qualified as Mild group and 5 and 6 as severe group.

Subjects were all informed and did not oppose, written consent for participation was not required for this study in accordance with the national legislation and the institutional requirements. The study is registered on the Health Data Hub website under the number F20210218154851. A commitment to comply with Reference Methodology n°004 issued by French Authorities (CNIL) has been signed by the investigator (Prof. O. Epaulard co-author of the current study). Anonymized patient data (age, sex, severity, body mass index) are described in the Supplementary File 2. Additional raw data regarding the cohort can be made available by the authors of the current study within respect of General Data Protection Regulation, without undue reservation.

#### Plasma sample pre-processing

Plasma samples had to be pre-processed to sustain *E. coli* growth, following the protocol described in Ahavi *et al.* [38]. Briefly, samples were deproteinized by centrifugation (1h, 4000g) in Amicon Ultra-4 Centrifugal Filters (3 kDa MWCO, 4 mL of sample volume), resuspended and re-centrifuged (1h, 4000g) in the same centrifugal filters. Filtrates were aliquoted and frozen at -80°C for storage. A 50 *µ*L aliquot of each patient sample was tested for residual antibiotics. This was done by growing *E. coli* on the sample according to the protocol described in the Growth assays on plasma subsection. Patient samples failing this test were excluded from the analysis.

#### Growth assays on plasma

Growth assays followed the protocol described in Ahavi *et al.* [38]. For a given mutant, the final test volume was 100 *µ*L. In a non-peripheral well of a clear flat-bottom 96-well plate, 1.02 *µ*mol of MgSO_4_, 4.78 *µ*mol of Na_2_HPO_4_, and 2.2 *µ*mol of KH_2_PO_4_ were added to deproteinized plasma (50, 25 or 12.5 *µ*L depending on the desired plasma dilution) to compensate for the EDTA-chelation of divalent cations and buffer the pH. 20 *µ*L of the 5X corresponding nutrient supplement (cf. Supplementary File 1) was also added to the well, which was then inoculated with a mutant culture prepared as follows. Sterile Milli-Q water was added to a final volume of 100 *µ*L.

The mutant glycerol stock was first recovered on a M9 plate supplemented with 0.07% D-(+)-glucose (to emulate plasmatic glucose concentration) and 0.5% yeast extract in order to adapt the strains for growth in plasma samples. The plate was incubated overnight at 37°C. An isolated colony was subsequently inoculated in 3mL of the corresponding liquid medium and incubated at 37°C until the *OD*600 reached between 0.5 and 1. This culture is then inoculated to the well at a final *OD*6*OO* of 0.005.

The plate was then inoculated in a BioTek Synergy HTX Multimode Reader (Agilent) at 37°C with continuous orbital shaking at 807cpm (1mm). Growth was assayed by *OD*600 monitoring every 10 minutes over periods ranging from 8h and 33h15min, depending on the mutant considered.

#### Sensor screening

All 34 mutants were screened following the protocol described in the Growth assays on plasma subsection.

Due to limitations regarding the number of patients and the volume of plasma per sample, sensor screening was performed on three biological replicates for each category (Mild, Severe, Negative) in order to preserve enough patient samples for the sensor validation phase. Since the volume of a pre-processed sample was insufficient to screen every mutant, plasma sample pools were prepared for each biological replicate. For each category, the plasma pool for replicate 1 included samples from eight patients, the plasma pool for replicate 2 included samples from four patients, and the plasma pool for replicate 3 included samples from four patients.

To maximize the likelihood that that the plasmatic concentration of each targeted metabolite across all three categories would fall within the dynamic range of its corresponding mutant, a plasma dilution selection step was performed during replicate one. Each mutant was tested sequentially on three plasma dilutions: 2-fold, 4-fold and 8-fold, until growth phenotype alterations became visible compared to a wild-type strain growing on plasma at the same dilution supplemented with all proteinogenic amino acids and nucleobases (the concentration of each supplement is indicated in Supplementary File 1 and inspired from Neidhardt *et al.* [64]. Replicates 2 and 3 were then performed at the selected dilution for each mutant that didn’t display uninterpretable growth profiles, residual growth, or no growth (Supplementary Figure 1).

#### Sensor validation

The six selected mutants were cultured in technical duplicate in every remaining patient sample, following the procedure detailed in the Growth assays on plasma subsection and using the plasma dilution selected during the sensor screening phase for each mutant. The validation sample set included 19 samples from COVID-negative subjects, 18 pre-onset samples from patients who later developed a mild form of the disease as defined in the Plasma sample collection subsection, and 19 pre-onset samples from patients who later developed a severe form of the disease as defined in the Plasma sample collection subsection. These samples were kept separate from the ones that were incorporated in the plasma pools used in the sensor screening phase from the outset of the study. The aim was to ensure a relative independence between both analyses and therefore strengthen result robustness, even though all patient samples came from the same cohort (detailed in the Plasma sample collection subsection).

### Statistical analyses of growth parameters

#### Growth parameter extraction

Baseline correction was applied to all growth curves analyzed in order to remove the contribution of the culture medium. Seven growth parameters were then extracted from the analyzed growth curves. Each of these parameters was chosen because biologically meaningful as they reflect underlying biological processes and vary depending on the physicochemical properties and nutrient richness of the culture medium.

*µ_max_*, the maximal growth rate, represents the highest rate of bacterial cell division. To estimate this parameter, a first time window covering the 6 to 8 time points with the highest local growth rates is computed from the derivative of the *ln*(*OD*600)*vstime* curve. A linear model of the form (*y* = *at* + *b*) was then fitted to these points. A mean squared error (MSE) is calculated between the observed and predicted values within the time window. Time points immediately upstream and downstream of the selected window were subsequently evaluated for inclusion. A candidate point was accepted only when its addition, followed by refitting of the linear model, resulted in a lower MSE. This procedure was repeated iteratively until adding points in either direction no longer improved the fit. The maximal growth rate was subsequently defined as the slope of the final linear model.

*OD*600*_max_* was defined in this study as the *OD*600 reached at beginning of the stationary phase. It served as a proxy for the maximal living biomass concentration. That value was collected at the beginning of the stationary phase because *OD*600 values were not yet substantially affected by the contribution of dead cells. Let *m_t_*denote the local slope of the raw *OD*600 values (without log transformation), *m*_max_ the mean of the five highest local slopes, and max(*OD*) the maximum recorded *OD*600 value. In this study, *OD*600_max_ was defined as the first OD600 value at time *t* satisfying:

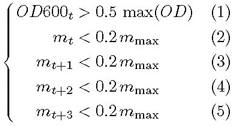

Equation (1) was added to ensure that no *OD*600 value from the lag phase could be selected. Regarding equations (2) to (5), 0.2 was set as an arbitrary threshold.

*ln*(*OD*600)*_max_* is the *ln*(*OD*600) value of the last point of the exponential phase. In this study, it served as a proxy for the biomass concentration at the end of the exponential phase (and before the deceleration phase when existing). In this study, the end of the exponential phase was defined as the first time point, *t*, after the time points used to calculate *µ*_max_, for which the following condition held:

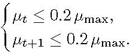

0.2 was set as an arbitrary threshold. *ln*(*OD*600)*_max_* was then defined as the *ln*(*OD*600) value at that time point.

The span of the lag phase reflects the adaptation time to the culture medium that the strain required before entering exponential phase. In this study, it was defined as the last time point, *t*, before the time points used to calculate *µ*_max_, for which the following condition held:

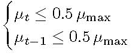

0.5 was set as an arbitrary threshold.

The span of the exponential phase is a parameter that is not redundant with *µ_max_*as *µ_max_*is usually inferred from a subset of points from the exponential phase corresponding to a growth-rate maximum. It also allows the span of the deceleration phase (when present) to be inferred. In cases where there is a deceleration phase, that is, this suboptimal growth phase sometimes following exponential phase, a gap will be observed between the end of the exponential phase and the beginning of the stationary phase. In this study, the span of the exponential phase is defined by the difference between the time point corresponding to *ln*(*OD*600)*_max_* and the end of the lag phase.

The start of the stationary phase corresponds to the time where the strain cannot grow exponentially anymore, due to nutrient depletion or unfavorable physicochemical conditions (such as low pH or toxin accumulation). It is defined in this study as the time point corresponding to the *OD*600*_max_* value.

The *OD*600*_max_* − *OD*600*_final_* difference is a proxy for the balance between growth and death during stationary phase. Mathematically, it is the difference between the *OD*600*_max_* and the *OD*600 value of the last time point recorded. A difference was preferred over a slope because stationary phases are not always linear.

All durations and time points are expressed in hours (*h*) and *µ_max_* is expressed in *h^−^*^1^.

#### Welch’s *t*-test on growth parameters

All growth parameters determined during sensor screening are provided Supplementary Data 2. Two-sided Welch’s *t*-tests [44] were performed to assess whether growth parameters are on average significantly different between the Mild and Severe groups, and between the Negative and Positive groups. Patient-level values (experimental units) were obtained by averaging the values of the parameters extracted from the two technical replicates. For each growth parameter, the null hypothesis was that its mean value was equal between the two groups being compared. Welch’s *t*-test was used because of its robustness against unequal variances and unbalanced sample sizes. A Benjamini–Hochberg [42] multiple-testing *p*-value correction was applied to account for the multiple-testing across the six strains and the seven growth parameters.

Welch’s *t*-test was conducted in Python using scipy.stats.ttest_ind [65] (v.1.11.1) (equal_var=False). statsmodels.stats.multitest [66] (v.0.14.6) was used to perform Benjamini–Hochberg [42] multiple-testing corrections. Statistical significance was assessed at *α* = 0.1. Means, sample sizes, degrees of freedom, t-statistics, and *p*-values are provided in Supplementary Data 2.

In the case of the Mild vs Severe vs Negative problem, statistical significance was assessed through pairwise comparisons between all three groups. An additional Benjamini–Hochberg [42] multiple-testing *p*-value correction was applied to account for the three pairwise comparisons.

### Statistic modeling of growth curves using Generalized Additive Mixed Models (GAMMs)

statsmodels.stats.multitest (v.0.14.6) was used to perform all Benjamini–Hochberg [42] multiple-testing corrections in the following subsections.

#### Growth curves from plasma sample pools

Baseline correction was applied to all growth curves analyzed in order to remove the contribution of the culture medium. All baseline-corrected growth curves acquired during sensor screening are provided in Supplementary Data 1 and in Supplementary Figures 13 to 18. Growth curves obtained from plasma pools during sensor screening were analyzed using generalized additive mixed models [41] (GAMMs) to identify strains whose growth phenotype varied in average according to sample group (mild, severe, negative).

Models were fitted in R using the bam() function from the mgcv package (v.1.9.3) [67]. Three candidate models of increasing complexity were considered. Let *k* denote the basis dimension of the global smooth and *l* the basis dimension of the curve-specific smooth in model 3.

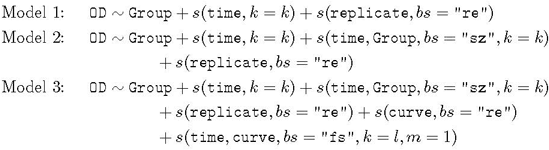

In all models, Group was included as a fixed effect to allow baseline differences between experimental conditions. The term s(time, k = k) modeled the overall nonlinear time trend. The term s(replicate, bs = “re”) represented a random intercept at the replicate level (across groups). The difference between the three models lay in the extent to which group-specific and curve-specific temporal deviations were allowed. No temporal autocorrelation correction within curves was applied.

Model 1 assumed a common temporal trajectory across groups, allowing only baseline shifts at the group level and at the replicate level. Model 2 added the term s(time, Group, bs = “sz”, k = k), allowing groups to differ in their temporal trajectories in addition to baseline differences. Model 3 further added s(curve, bs = “re”) and s(time, curve, bs = “fs”, k = l, m = 1), allowing each individual curve, defined as a replicate within a group, to have its own baseline level and smooth temporal deviation around the group-level trajectory.

For each classification problem (Mild vs Severe and Negative vs Positive), the three candidate models were fitted and compared using the Akaike Information Criterion (AIC), and the model with the lowest AIC was retained as the best-supported model among the candidates. The basis dimension *k* and, for model 3, the basis dimension *l*, were also optimized.

To select *k*, models with increasing values from 3 to 20 were fitted, and k.check() was used to assess adequacy of the basis dimension. The selected *k* was the smallest value yielding a k-index of at least 0.9. For model 3, *l* was explored from 3 to 20 after k was fixed, and the value yielding the lowest AIC was selected. The additional constraint *l* ≤ 150 × *m* was imposed.

For each mutant and each classification problem, the selected model was then fitted to the experimental data and approximate term-specific significance tests were extracted from the model summary. The global test is associated with the term s(time, Group, bs = “sz”, k = k) and the null hypothesis was that there was no group-specific difference in temporal trajectory. For the other smooth terms, the null hypothesis was that the corresponding term did not explain additional variability in the data beyond the rest of the model. Because each group included only three biological replicates owing to plasma volume restrictions, this stage of the analysis was treated as exploratory. Accordingly, the type I error rate was set at *α* = 0.1 and *p*-values were considered cautiously. To account for the multiple-testing across strains, *p*-values were corrected using the Benjamini-Hochberg procedure [42]. As the number of observations was too low to conclude about significance, an exclusion strategy was adopted, to filter out strains for which no evidence of group-related differences was detected.

In order to assess at which time windows the temporal trajectories shifted between groups, pairwise difference curves were computed from fitted GAMM predictions on a grid of 300 equally spaced time points spanning the analyzed time range. Within each group, a pointwise standard deviation of model-predicted values across replicate/curve units were computed. Moreover, because significance was assessed across the whole time course rather than in a pointwise manner, simultaneous confidence bands were generated instead of pointwise confidence intervals. Simultaneous 95% confidence bands were constructed for each difference curve using simulation from the estimated covariance matrix of the model coefficients. Specifically, 10,000 coefficient vectors were sampled from the approximate multivariate normal distribution of the fitted parameters, and for each simulated pairwise difference curve, the maximum absolute standardized deviation over the full time grid was computed. The 95th percentile of these maxima was then used as the simultaneous critical value for band construction. The same simulations were also used to conduct another global test of the null hypothesis of no difference between the two groups of curves over the entire time course. For this test also, a Benjamini-Hochberg [42] multiple-testing *p*-value correction was applied to account for multiple-testing across strains. For each classification problem, only strains that passed the two global tests described above (indicating that they were exhibiting group-specific phenotypic differences) were further examined to determine when these differences occurred. Time windows illustrating these differences were computed by merging adjacent time points at which the simultaneous confidence bands excluded 0.

In the case of the multiclass classification Mild vs Severe vs Negative, the term s(time, Group, bs = “sz”, k = k) tested the null hypothesis that all three groups followed the same temporal trajectory. It was therefore followed by three pairwise tests (Mild vs Severe, Mild vs Negative, and Severe vs Negative) using models whose basis dimensions had been optimized according to the procedure described earlier. An additional Benjamini–Hochberg [42] multiple-testing *p*-value correction was applied to account for the three pairwise comparisons. Accordingly, the time windows illustrating the periods when each group differed from one another were only computed for strains having passed the two global tests reported above and all three pairwise tests, indicating a significant difference between each group pair.

#### Growth curves from individual patients

All baseline-corrected growth curves acquired during sensor validation are provided in Supplementary Data 2. Because all remaining patient samples were included in this analysis, with 18, 19, and 19 patients in the Mild, Severe, and Negative groups, respectively, this GAMM analysis had greater statistical power than the initial mutant-screening step and was therefore used to assess significance. The same general strategy as in the mutant-screening phase was retained, although the candidate models differed because the experimental design was not the same. Let *k* denote the basis dimension of the global smooth, *l* the basis dimension of the patient-specific smooth, and *n* the basis dimension of the curve-specific smooth.

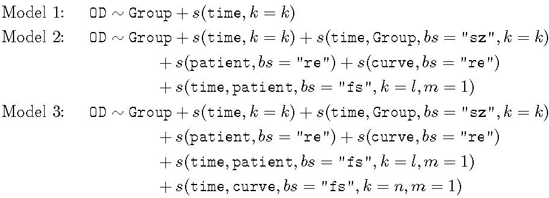

In all models, Group was included as a fixed effect in order to allow baseline differences experimental conditions, and *s*(time*, k* = *k*) was used to model the overall nonlinear time trend. No temporal autocorrelation correction within curves was applied. Model 1 assumed a common temporal trajectory across groups and only included baseline differences between groups. Model 2 additionally allowed group-specific temporal differences through the term s(time, Group, bs = “sz”, k = k). It also accounted for variability in baseline OD between patients and between curves through the random-intercept terms s(patient, bs = “re”) and s(curve, bs = “re”), where a curve denotes an individual longitudinal trajectory from one replicate of one patient. Finally, it allowed each patient to have its own smooth temporal deviation around the group-level trajectory through s(time, patient, bs = “fs”, k = l, m = 1). Model 3 extended this structure by also allowing each individual curve to have its own smooth temporal deviation through s(time, curve, bs = “fs”, k = n, m = 1).

As in the mutant-screening analysis, basis dimensions were optimized before final model comparison. The global basis dimension *k* was explored from 3 to 20 and assessed using k.check(). The selected value was the smallest one considered adequate according to the predefined k-index criterion. After *k* had been fixed, *l* and, for Model 3, *n* were explored from 3 to 20 and selected using AIC. Thus, *l* was optimized after fixing *k*, and *n* was optimized after fixing both *k* and *l*.

For each strain and each classification problem (Mild vs Severe, Negative vs Positive, and the three-group comparison), the selected model was then fitted to the corresponding dataset and approximate term-specific significance tests were extracted from the model summary. For the term s(time, Group, bs = “sz”, k = k), the null hypothesis was that there was no group-specific difference in temporal trajectory. For the other smooth terms, the null hypothesis was that the corresponding term did not explain additional variability in the data beyond the rest of the model. The type I error rate was set at *α* = 0.1 and *p*-values were corrected using a Benjamini-Hochberg [42] multiple-testing procedure to account for multiple-testing across the six strains.

Within each group, pointwise standard deviation of model-predicted values across replicate/curve units and simultaneous 95% confidence bands were computed as described in the Growth curves from plasma sample pools subsection. For each classification problem, only strains passing the two global tests described above (indicating that they were exhibiting group-specific phenotypic differences) were further examined to determine when these differences occurred. Time windows illustrating these differences were computed following the procedure described in the Growth curves from plasma sample pools subsection.

The case of the multiclass classification problem Mild vs Severe vs Negative was also handled following the same procedure described in the Growth curves from plasma sample pools subsection. Briefly, to ensure that each strain exhibited phenotypic differences allowing to differentiate each group from one another, all pairwise tests for the term the term s(time, Group, bs = “sz”, k = k) were conducted and an additional Benjamini–Hochberg [42] multiple-testing *p*-value correction was applied to account for the three pairwise comparisons. Moreover, the time windows corresponding to periods when each group differed from one another were only computed for strains having passed the two global tests (s(time, Group, bs = “sz”, k = k) term and simulation-based test) as well as all three pairwise tests.

### Classification model implementations

#### General architecture and metrics

All classifiers developed in this study followed a general architecture designed to reduce data leakage and provide robust performance estimates. First, model evaluation relied on a nested cross-validation framework comprising 5 outer folds and 3 inner folds.

Training was performed on each replicate but model evaluation was conducted at the patient-level to better approximate the real-world use case scenario and reduce biases arising from intra-patient correlations. Balanced accuracy was chosen over pooled outer-test area under the receiver operating characteristic curve (AUC) as the hyperparameter optimization objective mainly because final classification performance (class assignment) was preferred over probability ranking performance. A secondary reason is that the negative and positive classes in the diagnostic classification problem are imbalanced.

Regarding ensemble models, an out-of-fold (OOF) procedure was performed during ensemble weight optimization to avoid data leakage. Within each outer-training fold, patients were further split into OOF-training and OOF-validation partitions. Each base model was hyperparameter-tuned and trained using the corresponding OOF-training partition, while ensemble weight optimization was performed using predictions generated on the held-out OOF-validation partition. Replicate-level predictions were aggregated at the patient level prior to ensemble optimization.

Outer folds were stratified using StratifiedKFold (scikit-learn v.1.3.0 [68]), and patient-level separation was enforced so that each patient appeared only once in the test set, ensuring robust classification estimates and preventing biases arising from intra-patient correlations. For each classification task, a single fold partition was generated and reused across all models to enable fair model comparisons. In each case, the reported metrics include the mean, standard deviation, median, 2.5^th^ and 97.5^th^ percentiles for: balanced accuracy, macro precision, macro recall, macro F1-score, macro specificity, AUC across fold. Pooled outer-test AUC is also indicated. Metrics were computed using scikit-learn functions (v.1.3.0) [68]. Performance metrics associated with all classifiers, as well as hyperparameters, patient predictions, raw folds, ROC (receiver operating characteristic) curves and confusion matrices are available in a Zenodo repository [57] (DOI: 10.5281/zenodo.20585116). A summary report of the best-performing strain models for each classification approach across all strains and classification tasks is available in Supplementary Data 3.

#### Growth-parameter-based classification

Four classifiers were implemented to perform classical growth-curve classification based on the extracted growth parameters: (i) a support vector machine [69] (SVM) using SVC from scikit-learn (v.1.3.0) [68], (ii) a logistic regression model [70] using LogisticRegression from scikit-learn (v.1.3.0) [68], (iii) a gradient-boosted tree model XGBoost [71] using XGBClassifier from xgboost (v.3.2.0), and (iv) a soft-voting ensemble [72] combining the three base classifiers using VotingClassifier from scikit-learn (v.1.3.0) [68]. In order to reduce the risk of data leakage and ensure a robust performance assessment, a nested cross-validation strategy with 5 outer folds and 3 inner folds was implemented. Training of all models was performed after normalization of the input features using StandardScaler (scikit-learn v.1.3.0 [68]. Hyperparameters were optimized in the inner cross-validation loop using balanced accuracy as the selection criterion (Table 2), and the best configuration was re-fitted on the outer-training set.

**Table 2:** Hyperparameters tuned in the inner cross-validation loop for each classifier.

| Model | Hyperparameter | Search values |
| --- | --- | --- |
| SVM | $C$ | {0.1, 1, 10} |
| SVM | Kernel | {rbf} |
| Logistic regression | $C$ | {0.01, 0.1, 1, 10} |
| XGBoost | n_estimators | {100, 300} |
| XGBoost | max_depth | {2, 3, 5} |
| XGBoost | learning_rate | {0.03, 0.1} |
| Voting ensemble | Weights (SVM, logistic regression, XGBoost) | {1, 2, 3}<sup>3</sup> |

Model training was performed at the replicate level. For patient-level evaluation, replicate-level predicted probabilities were averaged across the two replicates of each patient, and the final class assignment was defined as the class with the highest averaged probability.

Regarding the soft-voting ensemble, an out-of-fold procedure was used during weight optimization to avoid information leakage. Each base classifier was re-tuned and re-fitted within the training folds, and the ensemble weights were selected using grid search exclusively from out-of-fold predictions generated on the outer-training set. A fixed random seed of 2026 was used to ensure reproducibility.

A univariate feature-selection procedure (SelectKBest and f_classif for the ANOVA F-test, both from scikit-learn [68] v.1.3.0) was also implemented but was not retained in the final analysis because it did not improve balanced accuracy (Supplementary Figure 3). When enabled, feature selection was applied exclusively within the training folds to prevent data leakage.

#### Time-series classification

In order to challenge the traditional growth-parameter-based classification framework, a full growth curve analysis was performed by feeding complete time series into two deep-learning models: a one-dimensional convolutional neural network [73] (CNN1D) and a temporal convolutional network [74] (TCN). The two models were implemented using PyTorch [75]. Here again, a nested cross-validation strategy with 5 outer folds and 3 inner folds was implemented to prevent data leakage. *z* − *score* normalization was applied to raw time series from the A1, A19 and A28 strains only and not to the ones from the A5 and A15 strains, as it might hide information about *OD*600 amplitude. Baseline-correction was applied to all growth curves analyzed in order to remove the contribution of the culture medium. Normalization parameters were estimated exclusively from the training partition within each fold and then applied to the corresponding validation or test partition (inner or outer loop) to avoid data leakage.

Hyperparameter optimization was performed using Optuna (v.4.7.0) [76] within the inner loop of the nested patient-level cross-validation procedure. For each trial, Optuna (v.4.7.0) [76] sampled a candidate hyperparameter configuration (Table 3) and its performance was evaluated by the mean patient-level balanced accuracy across the inner folds. The hyperparameter configuration selected in the inner loop, including the number of training epochs, was then reused to retrain the model on the full outer-training set before evaluation on the corresponding outer-test fold. For each patient, predictions were generated for both replicate curves, logits were converted to class probabilities using the softmax function, and the resulting probabilities were averaged across replicates to obtain a single patient-level prediction.

**Table 3:**
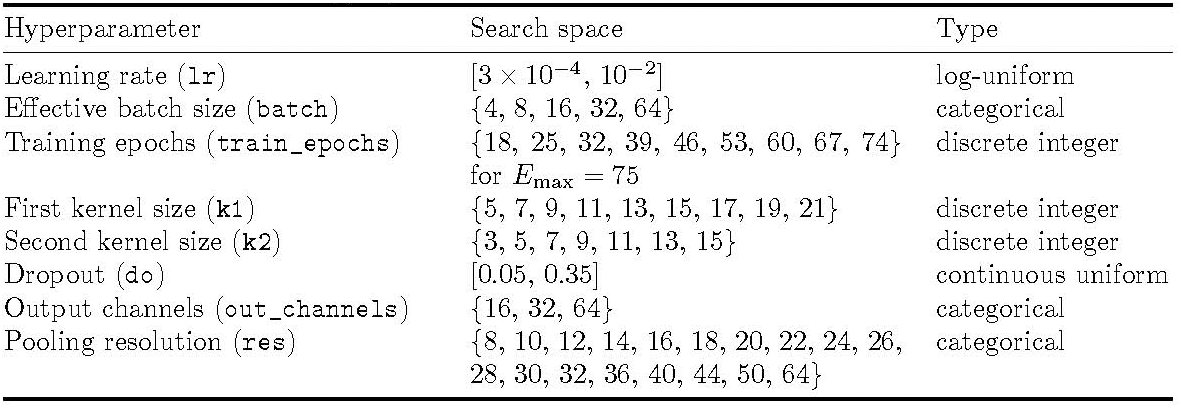
Hyperparameters tuned for the CNN and TCN models Hyperparameter Search space Type.

For the TCN, the dilation factor of the first temporal convolution block was additionally tuned with candidate values {1, 2, 4, 8}. The second TCN block used a fixed dilation of 1. After sampling, hyperparameters were further constrained to remain compatible with the input sequence length. In particular, pooling resolution was capped according to sequence length, and for the TCN the dilation could be reduced when the resulting receptive field became too large relative to the temporal dimension.

Optuna (v.4.7.0) [76] used a Tree-structured Parzen Estimator sampler with maximization of the inner-fold mean balanced accuracy as the optimization objective. No pruning strategy was applied. 60 trials were evaluated per outer fold.

Ray (v.2.54.0) [77] was used to execute independent outer-fold runs in parallel.

The first temporal derivative of the time series were also added as additional input channels to emphasize changes in slope and identify breakpoints. Channel-wise *z*−*score* normalization was applied to the derivatives of the time series from the A1, A19 and A28 strains only. Here also, normalization parameters were estimated exclusively from the training partition within each fold and then applied to the corresponding validation or test partition (inner loop or outer loop) to avoid data leakage.

#### Growth-parameter-conditioned time-series classification using Feature-wise Linear Modulation (FiLM)

In order to assess the information complementarity between the growth parameters and the features extracted from the growth curves by the deep-learning models and therefore increase the classification performances of each strain, the seven growth parameters extracted from each time series were added as contextual inputs within a feature-wise linear modulation (FiLM) framework [43]. In these multimodal architectures, the raw growth curve and its first temporal derivative were processed by either a CNN- and a TCN-based encoder to produce intermediate temporal feature maps. In parallel, the normalized growth parameters were passed through a multilayer perceptron that generated the FiLM parameters *γ* and *β*, which were then used to modulate the curve-derived feature maps through an affine transformation of the form *y* = *γ* ⊙*x*+*β*. Channel-wise *z* − *score* normalization was applied for the time series and the corresponding temporal derivatives of the A1, A19 and A28 strains only, following the same strategy implemented for time-series classification. For the A5 and A28 strains, the temporal derivative was not included as an input feature as its inclusion reduced the balanced accuracy of the CNN and TCN models during the time-series classification.

The FiLM parameters were generated by a dedicated multilayer perceptron applied to the normalized growth parameters. This FiLM generator comprised a linear layer, layer normalization, GELU activation, dropout and a final linear projection producing 2*C* outputs, where *C* denotes the number of channels in the temporal feature map. These outputs were split into scaling and shifting coefficients, with the scaling term parameterized as *γ* = 1 + *γ*_raw_, such that the modulation was initialized close to the identity transformation.

In addition, the final linear layer of the FiLM generator was initialized with zero weights and zero biases, yielding an initial condition of *γ* = 1 and *β* = 0. The hidden dimension of the FiLM generator was treated as a tunable hyperparameter during inner cross-validation (searched over 32, 64, 128).

In the present implementation, FiLM was applied according to a late-conditioning scheme. More specifically, modulation was not performed at the input level, but after the final convolutional or temporal encoding blocks, once high-level representations of the growth curves had already been extracted. The tabular branch therefore acted as a conditioning signal on these learned temporal features rather than on the raw signal itself.

In the CNN-based model, FiLM was applied after concatenation of the two parallel one-dimensional convolution branches. In the TCN-based model, FiLM was applied after the second temporal convolution block. In both cases, the modulated feature maps were then passed through adaptive average and max pooling layers, flattened, and fed to a final multilayer classifier.

Unlike feature-fusion approaches, this design allowed the growth parameters to directly condition the learned representation of the time series, thereby testing whether they could reweight the temporal features extracted from the growth curves.

#### Multistrain classification

A multistrain classifier was constructed in order to assess strain complementarity for each problem. Conceptually, each strain acts as a partial classifier, able to accurately classify a subset of patients. The goal was to assess whether combining several strains would allow patient-level information complementation of the models and therefore a better prognostic and diagnostic classification.

Two setups were implemented: (i) an early-fusion approach and (ii) a late-fusion approach. In the early-fusion configuration, the different strains were merged at the input level. The input configuration yielding the best balanced accuracy for each strain was included as additional channels and jointly fed to a CNN1D or a TCN. Model training then followed the procedure described in Time-series classification and Growth-parameter-conditioned time-series classification using Feature-wise Linear Modulation (FiLM) subsections. All combinations were tested among the 5 strains whose individual models showed classification ability for the Mild vs Severe problem (A28 was excluded).

Because the time series did not all have the same lengths, all time series involved were padded to the length of the longest time series before the normalization step. A binary mask was stored alongside each channel to distinguish valid observations from padded positions. This mask was propagated through the network, and mask-aware convolution, normalization (when required), and pooling operations were used so that padded regions did not contribute to feature extraction or aggregated representations.

The late fusion setup was a weighted soft-voting ensemble model. For each strain, the model and corresponding input feature set yielding the highest balanced accuracy in the strain-specific classification task were selected. Here also, all combinations were tested among the 5 strains whose individual models showed classification ability for the Mild vs Severe problem (A28 was excluded). Each base learner was first trained independently for its corresponding strain using the previously defined patient-level nested cross-validation workflow. The late-fusion step operated on the patient-level predicted class probabilities generated by these already trained base models.

Patient-level probabilities were formed by averaging the predicted probabilities of the 2 replicates within each patient. The final group prediction corresponded to the class with the highest mean probability. For each strain-specific model and for each outer fold, two types of probabilities were collected: (i) outer-test probabilities obtained from the held-out patients of the current outer fold and (ii) outer-train out-of-fold (OOF) probabilities obtained from patients belonging to the outer-training partition by cross-fitted prediction.

Fusion weights were learned independently within each outer fold using only the outer-training OOF probability table in order to prevent data leakage from the outer-test set. For each outer-fold, fusion performance on the outer-training set was first assessed by an additional internal cross-validation performed on this OOF probability table in order to assess the model performance in a robust manner. Weights were optimized using Optuna [76] (v.4.7.0), 300 trials per outer-fold) with balanced accuracy maximization as the optimization objective by searching one logit parameter per model in the interval [-6,6], followed by softmax transformation to obtain non-negative weights summing to 1.

Final fusion weights for a given outer fold were refit using the full outer-training OOF table. These final weights were then applied once to the aligned outer-test probabilities from the component strain-specific models to generate the fused outer-test predictions.

## Abbreviations

AUC: Area under the receiver operating characteristic curve
CNN1D: One-dimensional convolutional neural network
COVID-19: Coronavirus disease 2019
GAMM: Generalized additive mixed model
LogReg: Logistic regression
OD600: Optical density at 600nm
RT-PCR: Reverse transcription polymerase chain reaction
SVM: Support vector machine
TCN: Temporal convolutional network
XGBoost: Extreme Gradient Boosting algorithm

## Acknowledgements

The authors thank Candice Trocmé for assistance with plasma preparation, sample selection, and shipping. The authors acknowledge support from the French National Research Agency (ANR) under the grant ANR-21-CE45-0021-01 (AMN project). J.L.F. and P.A. also acknowledge funding provided by the ANR funding agency (grant number ANR-22-PEBB-0008 - PEPR B-BEST France 2030 program). J.L.F. additionally acknowledges support from the UE HORIZON BIOS program (grant number 101070281). A.L.G. is supported by the ANR funding agency (grant numbers ANR-21-CE45-0021-01 and ANR-24-CE44-1190), the University Grenoble Alpes Foundation as well as the Air Liquide Foundation (BIOMARCOVID Project).

## Supporting information

This article is associated with 2 supplementary files, 3 supplementary data compressed archives, 18 supplementary figures, one GitHub repository (https://github.com/brsynth/mutant-covid) and 2 Zenodo reposi-tories [56, 57], available free of charge.

- Supplementary Figures 1 to 18 are provided in a single file.
- Supplementary File 1 lists the generated mutants as well as the nutrient mixture and plasma dilution corresponding to each mutant.
- Supplementary File 2 details patient data (age, sex, severity, body mass index).
- Supplementary Data 1 and 2 compile all results (growth curves, growth parameters, GAMM analysis, statistical tests) acquired during sensor screening and sensor validation, respectively.
- Supplementary Data 3 provides summary reports of the best-performing models for each classification approach across all strains and classification tasks.
- All supplementary files and data are provided in a single file.
- Detailed classification results, including performance metrics, confusion matrices, patient predictions, hyperparameters, raw folds and ROC curves are available in a Zenodo repository [57] (DOI: 10.5281/zen-odo.20585116).
- All source codes (classifiers and GAMMs) are available in another Zenodo repository [56] (DOI: 10.5281/zenodo.20584758), as well as in this GitHub repository: https://github.com/brsynth/mutant-covid.

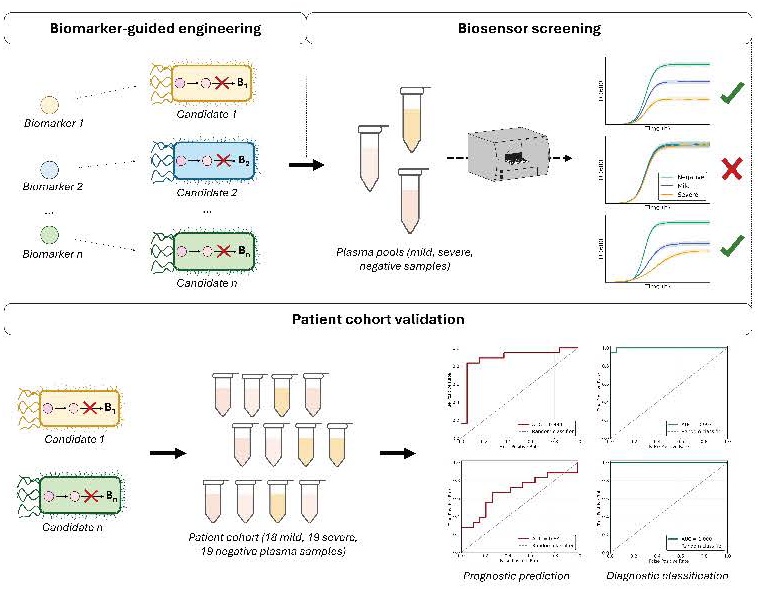

## References

(1) Chen, Y. et al. Nature Communications 2024, 15, 1657.

(2) Castelli, F. A.; Rosati, G.; Moguet, C.; Fuentes, C.; Marrugo-Ramírez, J.; Lefebvre, T.; Volland, H.; Merkoçi, A.; Simon, S.; Fenaille, F.; Junot, C. Analytical and Bioanalytical Chemistry 2022, 414, 759–789.

(3) Armitage, E. G.; Southam, A. D. Metabolomics 2016, 12, 146.

(4) Hocher, B.; Adamski, J. Nature Reviews Nephrology 2017, 13, 269–284.

(5) Ussher, J. R.; Elmariah, S.; Gerszten, R. E.; Dyck, J. R. Journal of the American College of Cardiology 2016, 68, 2850–2870.

(6) K. Trivedi, D.; A. Hollywood, K.; Goodacre, R. European Journal of Molecular & Clinical Medicine 2017, 3, 294.

(7) Rahman, M.; Schellhorn, H. E. Frontiers in Molecular Biosciences 2023, 10, 1120376.

(8) Jannetto, P. J.; Fitzgerald, R. L. Clinical Chemistry 2016, 62, 92–98.

(9) Saltepe, B.; Kehribar, E. Ş.; Su Yirmibeşoğlu, S. S.; Şafak Şeker, U. Ö. ACS Sensors 2018, 3, 13–26.

(10) Gao, Y.; Wang, L.; Wang, B. Nature Communications 2023, 14, 8415.

(11) Kotula, J. W.; Kerns, S. J.; Shaket, L. A.; Siraj, L.; Collins, J. J.; Way, J. C.; Silver, P. A. Proceedings of the National Academy of Sciences 2014, 111, 4838–4843.

(12) Brophy, J. A. N.; Voigt, C. A. Nature Methods 2014, 11, 508–520.

(13) Qin, L.; Liu, X.; Xu, K.; Li, C. Current Opinion in Biotechnology 2022, 75, 102694.

(14) Darmostuk, M.; Rimpelova, S.; Gbelcova, H.; Ruml, T. Biotechnology Advances 2015, 33, 1141–1161.

(15) Rizik, L.; Danial, L.; Habib, M.; Weiss, R.; Daniel, R. Nature Communications 2022, 13, 5602.

(16) Zhang, L.; Guo, W.; Lu, Y. Biotechnology Journal 2020, 15, 2000187.

(17) Schulz-Mirbach, H.; Dronsella, B.; Erb, T. J. Trends in Biotechnology 2026, 44, 92–110.

(18) Orsi, E. et al. Nature Communications 2025, 16, 2168.

(19) Jiang, W.; Bikard, D.; Cox, D.; Zhang, F.; Marraffini, L. A. Nature Biotechnology 2013, 31, 233–239.

(20) González-Delgado, A., et al. Nature Biotechnology 2026, DOI: 10.1038/s41587-026-03076-6.

(21) Monk, J. M.; Lloyd, C. J.; Brunk, E.; Mih, N.; Sastry, A.; King, Z.; Takeuchi, R.; Nomura, W.; Zhang, Z.; Mori, H.; Feist, A. M.; Palsson, B. O. Nature Biotechnology 2017, 35, 904–908.

(22) Lloyd, C. J.; King, Z. A.; Sandberg, T. E.; Hefner, Y.; Olson, C. A.; Phaneuf, P. V.; O’Brien, E. J.; Sanders, J. G.; Salido, R. A.; Sanders, K.; Brennan, C.; Humphrey, G.; Knight, R.; Feist, A. M. PLOS Computational Biology 2019, 15, ed. by Hatzimanikatis, V., e1006213.

(23) Sleight, S. C.; Bartley, B. A.; Lieviant, J. A.; Sauro, H. M. Journal of Biological Engineering 2010, 4, 12.

(24) Masson, J.-F. ACS Sensors 2020, 5, 3290–3292.

(25) Quispe Haro, J. J.; Wegner, S. V. Advanced Healthcare Materials 2023, 12, 2300835.

(26) Zúñiga, A.; Muñoz-Guamuro, G.; Boivineau, L.; Mayonove, P.; Conejero, I.; Pageaux, G.-P.; Altwegg, R.; Bonnet, J. Frontiers in Bioengineering and Biotechnology 2022, 10, 859600.

(27) Tepper, N.; Shlomi, T. PLoS ONE 2011, 6, ed. by Peccoud, J., e16274.

(28) Dong, H. et al. Biotechnology Advances 2026, 86, 108746.

(29) Hernández-Sancho, J. M.; Boudigou, A.; Alván-Vargas, M. V. G.; Freund, D.; Arnling Bååth, J.; Westh, P.; Jensen, K.; Noda-García, L.; Volke, D. C.; Nikel, P. I. Nature Communications 2024, 15, 8316.

(30) Thomas, B. et al. Emergency Medicine Journal 2021, 38, 587–593.

(31) Reynard, C.; Allen, J. A.; Shinkins, B.; Prestwich, G.; Goves, J.; Davies, K.; Body, R. Emergency Medicine Journal 2022, 39, 70–76.

(32) Shen, B. et al. Cell 2020, 182, 59–72.e15.

(33) Danlos, F.-X., et al. Cell Death & Disease 2021, 12, 258.

(34) Roberts, I.; Wright Muelas, M.; Taylor, J. M.; Davison, A. S.; Xu, Y.; Grixti, J. M.; Gotts, N.; Sorokin, A.; Goodacre, R.; Kell, D. B. Metabolomics 2022, 18, 6.

(35) Correia, B. S. B.; Ferreira, V. G.; Piagge, P. M. F. D.; Almeida, M. B.; Assunção, N. A.; Raimundo, J. R. S.; Fonseca, F. L. A.; Carrilho, E.; Cardoso, D. R. Journal of Proteome Research 2022, 21, 1640–1653.

(36) López-Hernández, Y.; Monárrez-Espino, J.; Oostdam, A.-S. H.-v.; Delgado, J. E. C.; Zhang, L.; Zheng, J.; Valdez, J. J. O.; Mandal, R.; González, F. D. L. O.; Moreno, J. C. B.; Trejo-Medinilla, F. M.; López, J. A.; Moreno, J. A. E.; Wishart, D. S. Scientific Reports 2021, 11, 14732.

(37) Sindelar, M.; Stancliffe, E.; Schwaiger-Haber, M.; Anbukumar, D. S.; Adkins-Travis, K.; Goss, C. W.; O’Halloran, J. A.; Mudd, P. A.; Liu, W.-C.; Albrecht, R. A.; García-Sastre, A.; Shriver, L. P.; Patti, G. J. Cell Reports Medicine 2021, 2, 100369.

(38) Ahavi, P.; Hoang, T.-N.-A.; Meyer, P.; Berthier, S.; Fiorini, F.; Castelli, F.; Epaulard, O.; Le Gouellec, A.; Faulon, J.-L. Cell Systems 2026, 101654.

(39) Meyer, F.; Keller, P.; Hartl, J.; Gröninger, O. G.; Kiefer, P.; Vorholt, J. A. Nature Communications 2018, 9, 1508.

(40) Gómez-Coronado, P. A.; Kubis, A.; Kowald, M.; Ute, R.; Cotton, C.; Lindner, S. N.; Bar-Even, A.; Erb, T. J. Synthetic Biology 2025, 10, ysaf004.

(41) Lin, X.; Zhang, D. Journal of the Royal Statistical Society Series B: Statistical Methodology 1999, 61, 381–400.

(42) Benjamini, Y.; Hochberg, Y. Journal of the Royal Statistical Society Series B: Statistical Methodology 1995, 57, 289–300.

(43) Perez, E.; Strub, F.; de Vries, H.; Dumoulin, V.; Courville, A. FiLM: Visual Reasoning with a General Conditioning Layer, Version Number: 2, 2017.

(44) Welch, B. L. Biometrika 1947, 34, 28–35.

(45) Green, D. A.; Zucker, J.; Westblade, L. F.; Whittier, S.; Rennert, H.; Velu, P.; Craney, A.; Cushing, M.; Liu, D.; Sobieszczyk, M. E.; Boehme, A. K.; Sepulveda, J. L. Journal of Clinical Microbiology 2020, 58, ed. by McAdam, A. J., e00995–20.

(46) Xia, J.; Broadhurst, D. I.; Wilson, M.; Wishart, D. S. Metabolomics 2013, 9, 280–299.

(47) Qiu, S.; Cai, Y.; Yao, H.; Lin, C.; Xie, Y.; Tang, S.; Zhang, A. Signal Transduction and Targeted Therapy 2023, 8, 132.

(48) Volke, D. C.; Orsi, E.; Nikel, P. I. Current Opinion in Microbiology 2023, 75, 102353.

(49) Hutchison, C. A., et al. Science 2016, 351, aad6253.

(50) Jiang, W.; Wu, Z.; Gao, Z.; Wan, M.; Zhou, M.; Mao, C.; Shen, J. ACS Nano 2022, 16, 15705–15733.

(51) Gaut, N. J.; Deich, C.; Cash, B.; Hoog, T.; Engelhart, A. E.; Adamala, K. P. A Chemically Defined Synthetic Cell Capable Of Growth And Replication, en, 2026.

(52) Tica, J.; Oliver Huidobro, M.; Zhu, T.; Wachter, G. K.; Pazuki, R. H.; Bazzoli, D. G.; Scholes, N. S.; Tonello, E.; Siebert, H.; Stumpf, M. P.; Endres, R. G.; Isalan, M. Cell Systems 2024, 15, 1123–1132.e3.

(53) Cover, T. M. IEEE Transactions on Electronic Computers 1965, EC-14, 326–334.

(54) Slomovic, S.; Pardee, K.; Collins, J. J. Proceedings of the National Academy of Sciences 2015, 112, 14429–14435.

(55) Liu, D.; Wang, J.; Wu, L.; Huang, Y.; Zhang, Y.; Zhu, M.; Wang, Y.; Zhu, Z.; Yang, C. TrAC Trends in Analytical Chemistry 2020, 122, 115701.

(56) Ahavi, P.; Hoang, T.-N.-A.; Meyer, P.; Berthier, S.; Epaulard, O.; Le Gouellec, A.; Faulon, J.-L. Source code for “Engineering growth-coupled metabolic biosensors for disease prognosis and diagnosis using full growth trajectories”, 10.5281/zenodo.20584758, 2026.

(57) Ahavi, P.; Hoang, T.-N.-A.; Meyer, P.; Berthier, S.; Epaulard, O.; Le Gouellec, A.; Faulon, J.-L. Supporting data for “Engineering growth-coupled metabolic biosensors for disease prognosis and diagnosis using full growth trajectories”, 10.5281/zenodo.20585116, 2026.

(58) Kanehisa, M. Nucleic Acids Research 2000, 28, 27–30.

(59) Karp, P. D. et al. EcoSal Plus 2025, 13, ed. by Lovett, S. T., eesp–0019–2024.

(60) Jiang, Y.; Chen, B.; Duan, C.; Sun, B.; Yang, J.; Yang, S. Applied and Environmental Microbiology 2015, 81, ed. by Kelly, R. M., 2506–2514.

(61) Salgado, H., et al. Nucleic Acids Research 2024, 52, D255–D264.

(62) The UniProt Consortium, et al. Nucleic Acids Research 2025, 53, D609–D617.

(63) Labun, K.; Montague, T. G.; Krause, M.; Torres Cleuren, Y. N.; Tjeldnes, H.; Valen, E. Nucleic Acids Research 2019, 47, W171–W174.

(64) Neidhardt, F. C.; Bloch, P. L.; Smith, D. F. Journal of Bacteriology 1974, 119, 736–747.

(65) Virtanen, P. et al. Nature Methods 2020, 17, 261–272.

(66) Seabold, S.; Perktold, J. In Proceedings of the 9th Python in Science Conference, Austin, Texas, 2010, pp 92–96.

(67) Wood, S. N.; Goude, Y.; Shaw, S. Journal of the Royal Statistical Society Series C: Applied Statistics 2015, 64, 139–155.

(68) Kramer, O. In Machine Learning for Evolution Strategies, Series Title: Studies in Big Data; Springer International Publishing: Cham, 2016; Vol. 20, pp 45–53.

(69) Cortes, C.; Vapnik, V. Machine Learning 1995, 20, 273–297.

(70) Hosmer, D. W.; Lemeshow, S.; Sturdivant, R. X., Applied logistic regression, John Wiley & Sons, 2013.

(71) Chen, T.; Guestrin, C. In Proceedings of the 22nd ACM SIGKDD International Conference on Knowledge Discovery and Data Mining, ACM: San Francisco California USA, 2016, pp 785–794.

(72) Zhou, Z.-H., Ensemble Methods: Foundations and Algorithms, 2nd ed.; Chapman and Hall/CRC: Boca Raton, 2025.

(73) Lecun, Y.; Bottou, L.; Bengio, Y.; Haffner, P. Proceedings of the IEEE 1998, 86, 2278–2324.

(74) Bai, S.; Kolter, J. Z.; Koltun, V. An Empirical Evaluation of Generic Convolutional and Recurrent Networks for Sequence Modeling, Version Number: 2, 2018.

(75) Imambi, S.; Prakash, K. B.; Kanagachidambaresan, G. R. In Programming with TensorFlow, Prakash, K. B., Kanagachidambaresan, G. R., Eds., Series Title: EAI/Springer Innovations in Communication and Computing; Springer International Publishing: Cham, 2021, pp 87–104.

(76) Akiba, T.; Sano, S.; Yanase, T.; Ohta, T.; Koyama, M. In Proceedings of the 25th ACM SIGKDD International Conference on Knowledge Discovery & Data Mining, ACM: Anchorage AK USA, 2019, pp 2623–2631.

(77) Moritz, P.; Nishihara, R.; Wang, S.; Tumanov, A.; Liaw, R.; Liang, E.; Elibol, M.; Yang, Z.; Paul, W.; Jordan, M. I.; Stoica, I. Ray: A Distributed Framework for Emerging AI Applications, Version Number: 2, 2017.

