## Supplementary Figures for "Engineering growth-coupled metabolic biosensors for disease prognosis and diagnosis using full growth trajectories"

Supplementary figures 1 to 18 are provided below.

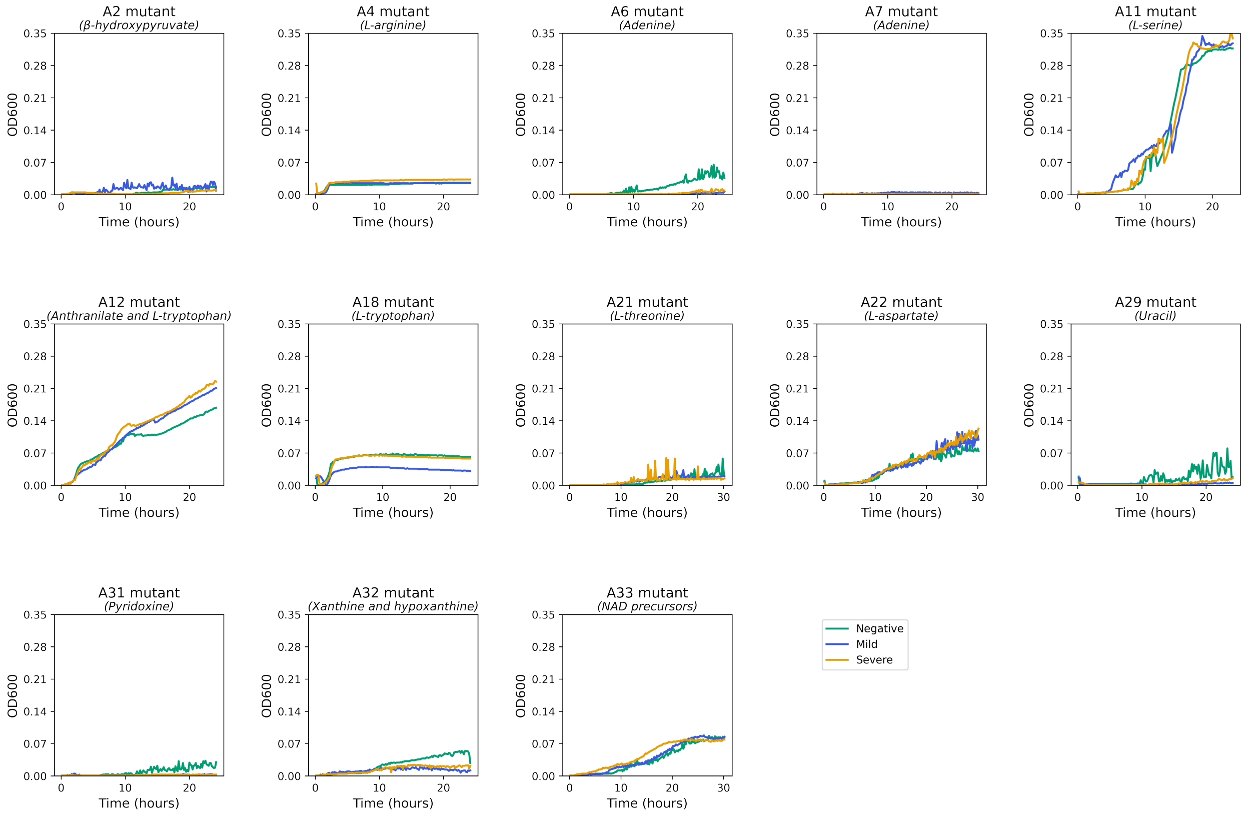


Supplementary Figure 1: Metabolic candidate sensors displaying inconclusive growth phenotypes.
This figure shows growth curves of mutants displaying residual or no growth, as well as mutants with uninterpretable growth profiles (A11, A12, A22, and A33 strains) on plasma samples.

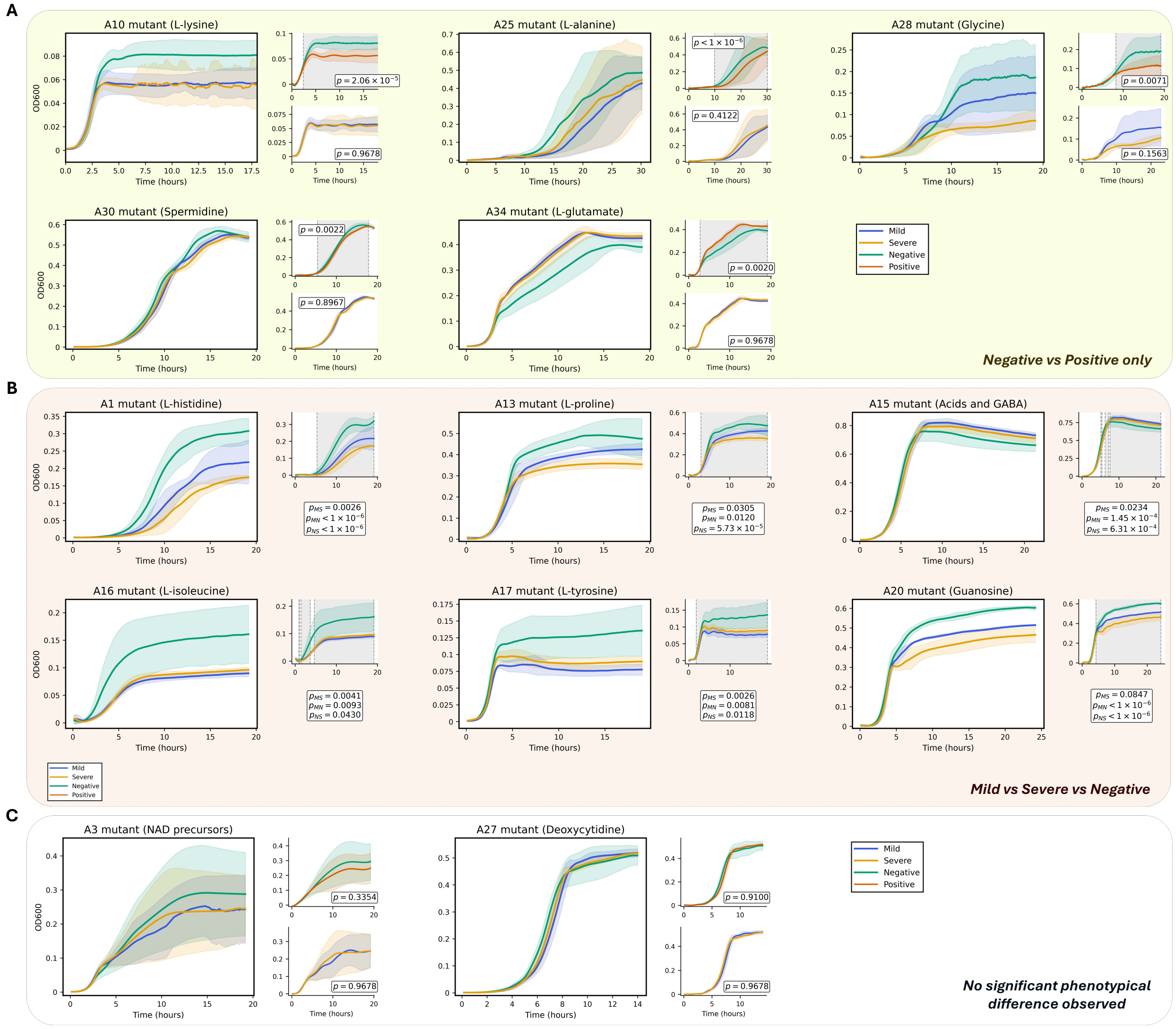


Supplementary Figure 2: Metabolic candidate sensors with COVID-19 diagnostic and multiclass classification potential.

Prognostic, diagnostic and multiclass classification potential was assessed using generalized additive mixed models [1] (GAMMs) applied to growth curves obtained from mutants cultured in mild, severe, and negative plasma sample pools ($n = 3$, $\alpha = 0.1$). Panel A shows mutants with diagnostic potential only. Panel B shows mutants with multiclass classification potential (mild vs severe vs negative). Panel C shows mutants without any identified prognostic or diagnostic potential. For each mutant, mean growth curves are indicated in the left section, GAMM-fitted mild and severe growth curves in the upper-right section and GAMM-fitted negative and positive growth curves in the lower-right section. Mild curves are colored in blue, severe curves in orange, negative curves in green and positive curves in red. Corresponding $p$-values (corrected using Benjamini-Hochberg procedure [2] to account for multiple testing) are indicated in each section. In panel B, "MS" stands for "mild vs severe", "MN" for "mild vs negative" and "NP" for "negative vs severe". Shaded areas represent standard deviations, and hatched areas indicate 95% confidence intervals (not readily visible in most plots owing to their narrow range). Grey windows in the GAMM representations indicate time intervals showing significant differences between the two conditions.

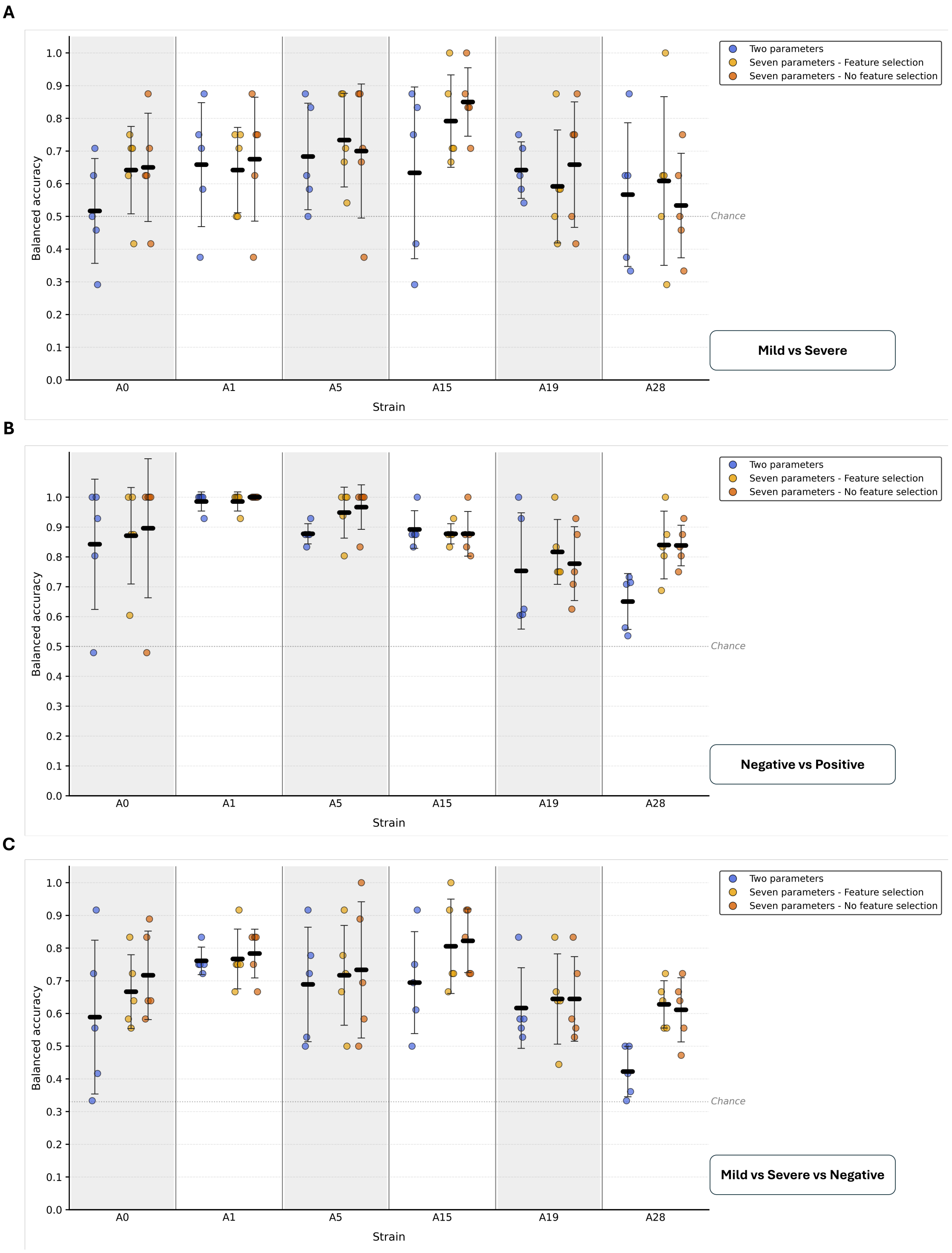


Supplementary Figure 3: Comparison of the classification performance using two and seven growth parameters as feature sets.

Classification performance is illustrated using balanced accuracy. Panels A, B, and C show, for *E. coli* K-12 MG1655 and the five selected metabolic candidate sensors, the highest balanced accuracy obtained for prognosis (mild vs severe classification), diagnosis (negative vs positive classification), and multiclass classification (mild vs severe vs negative), respectively, using machine learning classifiers (support vector machine, logistic regression, gradient-boosted trees and an ensemble model) trained on either the conventional two ($\mu_{max}$ and $OD600_{max}$) or the seven growth parameters identified in this study ($\mu_{max}$, $OD600_{max}$, $ln(OD600)_{max}$, lag phase duration, exponential phase duration, start of the stationary phase and $OD600_{max} - OD600_{final}$ difference). Results are shown with and without feature selection. Each dot represents one outer cross-validation fold ($n = 5$). The mean is indicated by a black bar, and error bars denote the standard deviation. The grey line at a balanced accuracy of 0.5 or 0.33 (for multiclass classification) indicates chance-level performance.

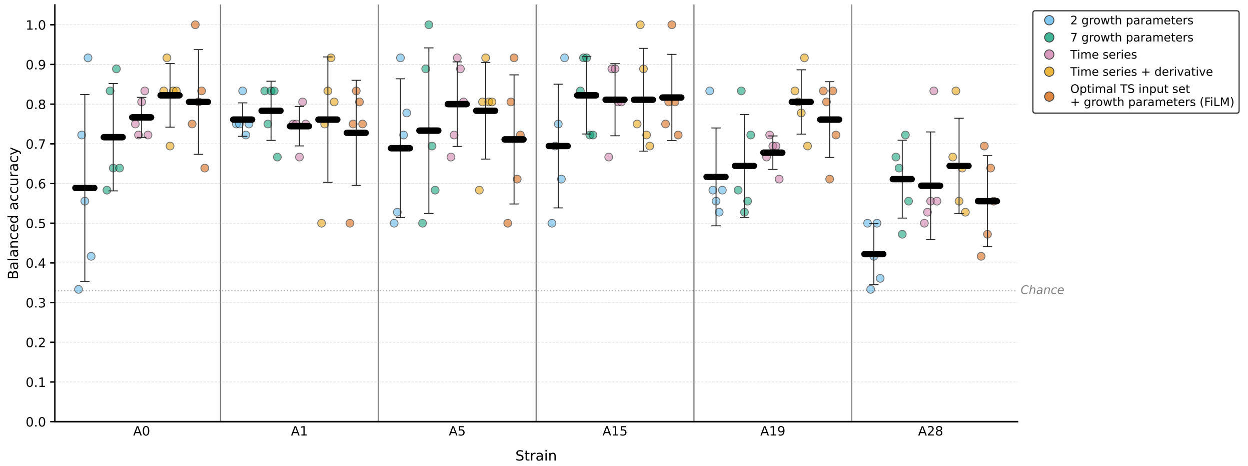


Supplementary Figure 4: Multiclass classification using the five selected metabolic sensors.
The figure shows, for the five selected metabolic candidate sensors, the highest balanced accuracy obtained for multiclass classification (mild vs severe vs negative) using different feature sets: (i) 2 growth parameters, (ii) 7 growth parameters, (iii) time series, (iv) time series combined with the first derivative, and (v) the best-performing time series input set combined with growth-parameter conditioning through feature-wise linear modulation [3] (FiLM). The grey line at a balanced accuracy of 0.33 indicates chance-level performance.

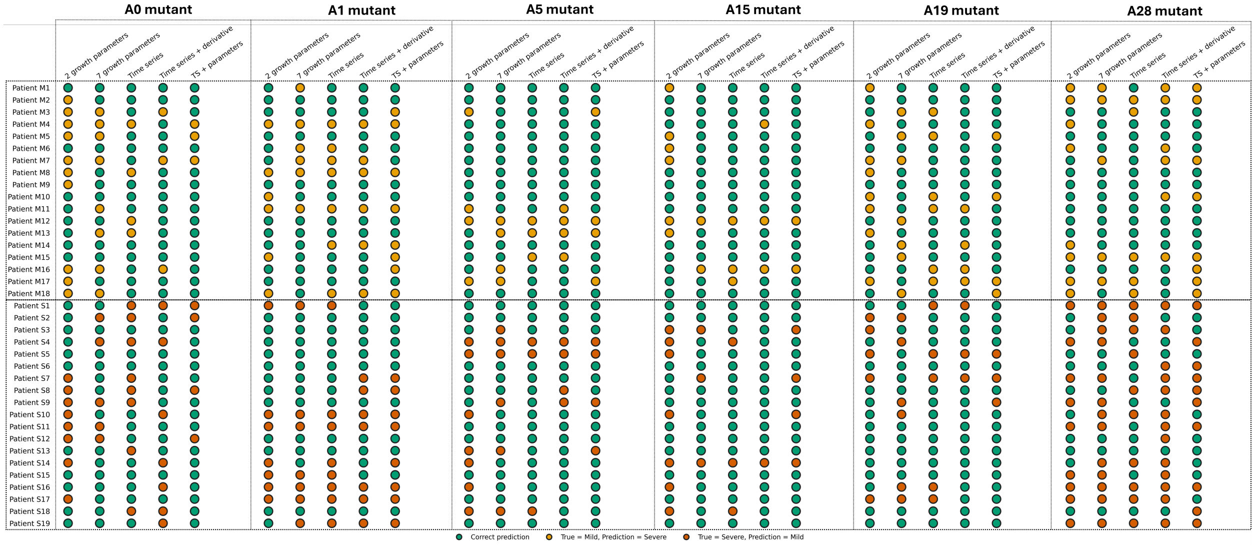


Supplementary Figure 5: Outer-fold cross-validation patient-level predictions for prognostic classification.
This figure shows, for each mutant, the outer-fold cross-validation patient-level predictions of the classifier achieving the highest balanced accuracy for prognostic classification using each feature set: (i) 2 growth parameters, (ii) 7 growth parameters, (iii) time series, (iv) time series combined with the first derivative, and (v) the best-performing time series input set combined with growth-parameter conditioning through feature-wise linear modulation (TS + parameters). Green dots indicate correct predictions, yellow dots indicate mild patients misclassified as severe, and orange dots indicate severe patients misclassified as mild.

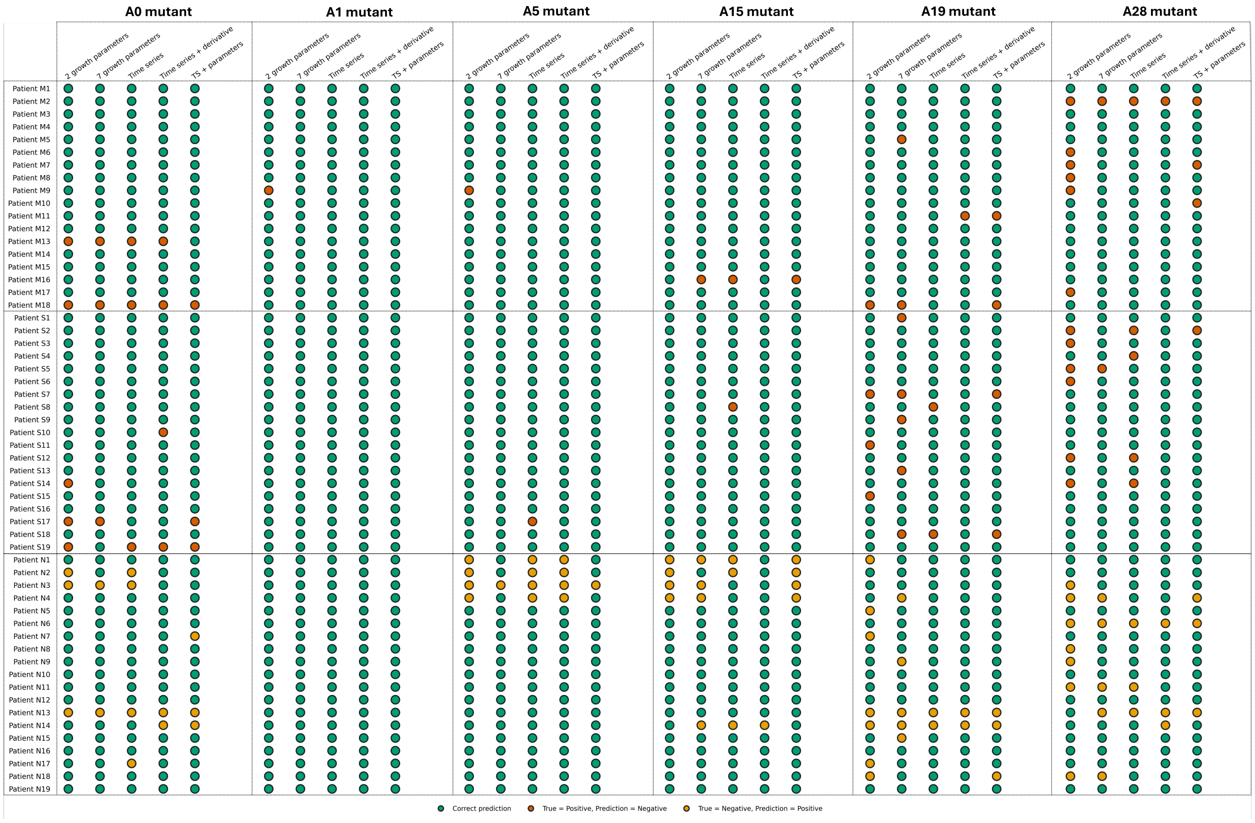


Supplementary Figure 6: Outer-fold cross-validation patient-level predictions for diagnostic classification. This figure shows, for each mutant, the outer-fold cross-validation patient-level predictions of the classifier achieving the highest balanced accuracy for diagnostic classification using each feature set: (i) 2 growth parameters, (ii) 7 growth parameters, (iii) time series, (iv) time series combined with the first derivative, and (v) the best-performing time series input set combined with growth-parameter conditioning through feature-wise linear modulation (TS + parameters). Green dots indicate correct predictions, orange dots indicate COVID-19 positive patients misclassified as COVID-19 negative individuals, and orange dots indicate COVID-19 negative individuals misclassified as COVID-19 positive patients.

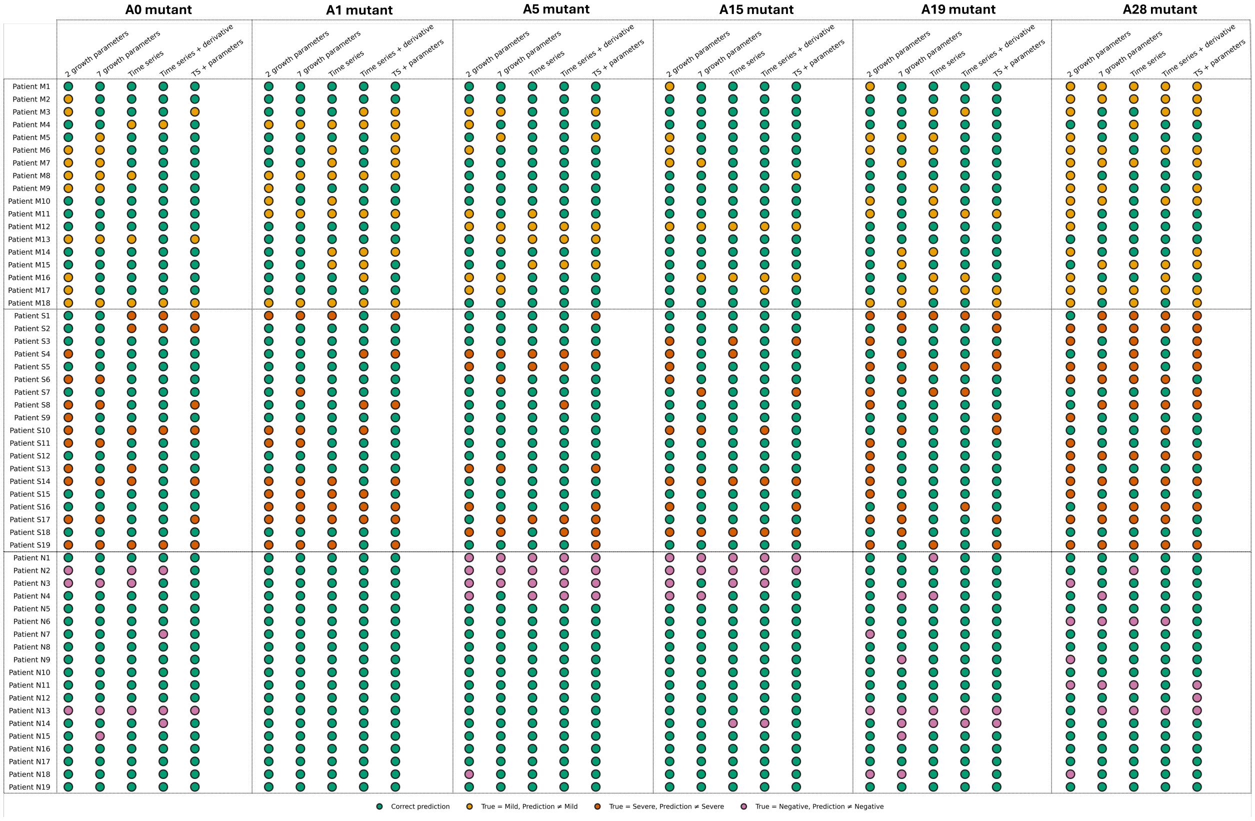


Supplementary Figure 7: Outer-fold cross-validation patient-level predictions for multiclass classification. This figure shows, for each mutant, the outer-fold cross-validation patient-level predictions of the classifier achieving the highest balanced accuracy for multiclass classification using each feature set: (i) 2 growth parameters, (ii) 7 growth parameters, (iii) time series, (iv) time series combined with the first derivative, and (v) the best-performing time series input set combined with growth-parameter conditioning through feature-wise linear modulation (TS + parameters). Green dots indicate correct predictions, yellow dots indicate misclassified mild patients, orange dots indicate misclassified severe patients and pink dots indicate misclassified negative subjects.

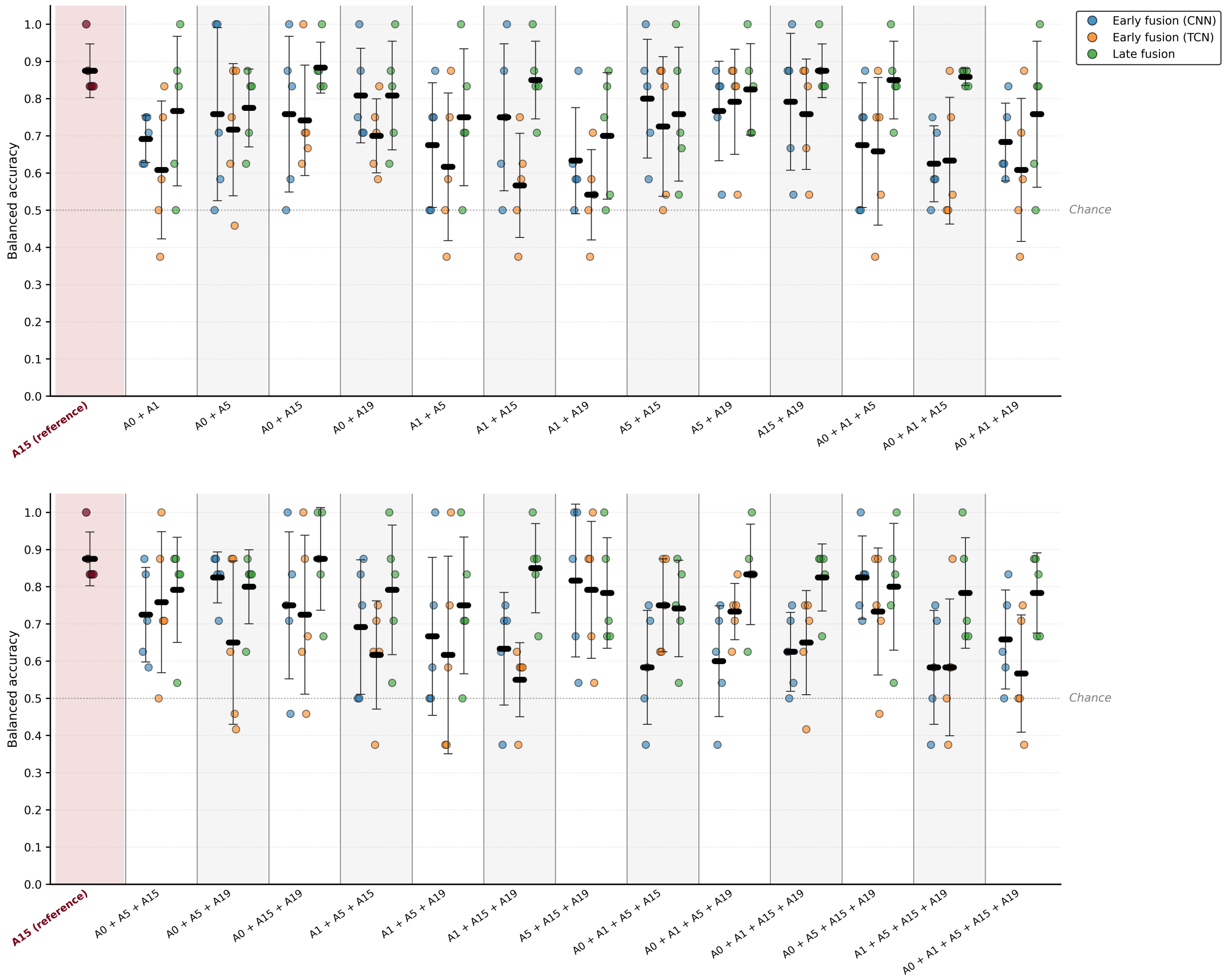


Supplementary Figure 8: Prognostic classification by multistrain models.

This figure shows the balanced accuracy of fusion models (early-fusion one-dimensional convolutional neural network, early-fusion temporal neural network and weighted soft-voting late-fusion ensemble model) trained on feature sets derived from all combinations of strains displaying prognostic capabilities (A0, A1, A5, A15 and A19). For each strain, the feature set used was the one associated with the highest balanced accuracy in the corresponding individual-strain classifier: (i) time series and first derivative (both not normalized) combined with growth-parameter conditioning through feature-wise linear modulation (FiLM) for A0, (ii) time series combined with the first derivative (both normalized) for A1, (iii) time series only (not normalized) for A5, (iv) time series combined with the first derivative (both not normalized) for A15 and (v) time series combined with the first derivative (both normalized) for A19. The individual-strain model yielding the highest balanced accuracy for prognosis, "A15 (reference)", is indicated for comparison.

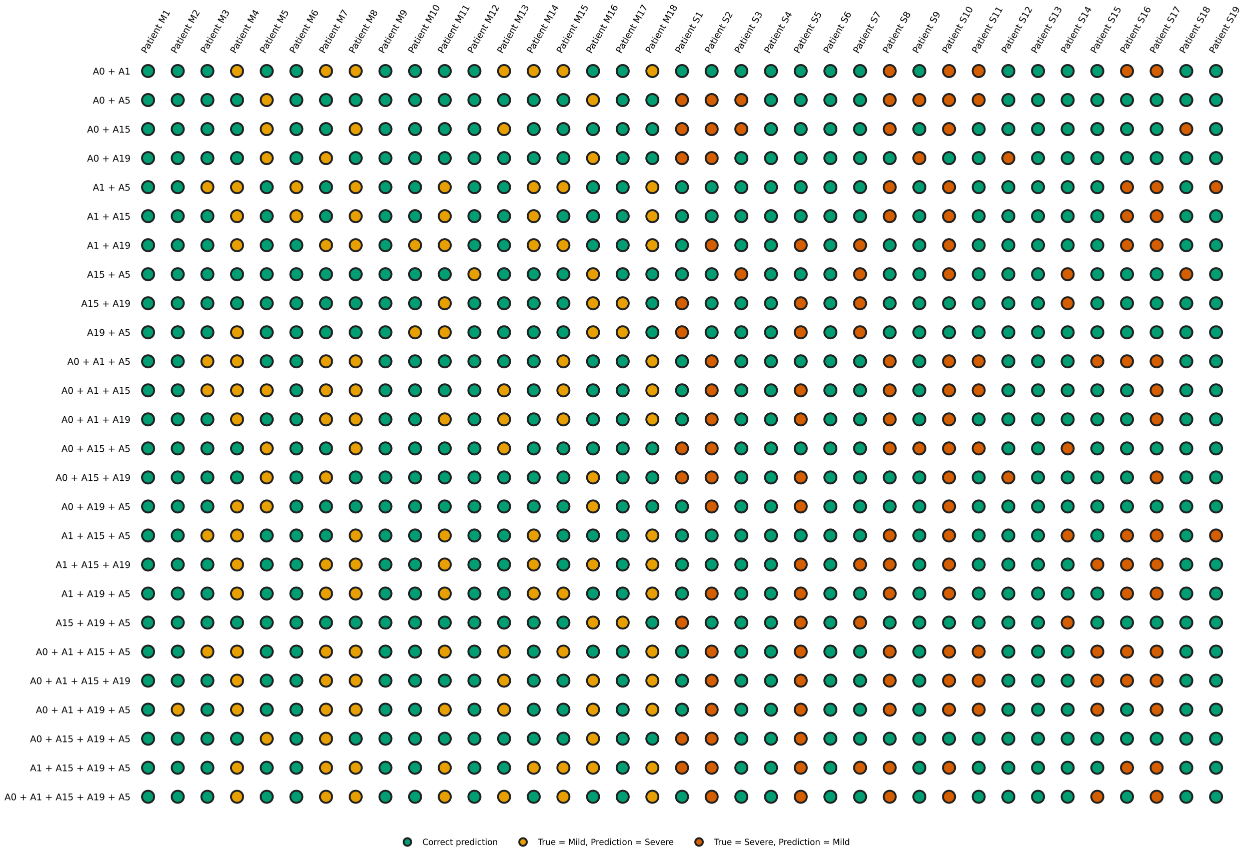


Supplementary Figure 9: Outer-fold cross-validation patient-level predictions of early-fusion one-dimensional convolutional neural network (CNN1D) models for prognostic classification.

Patient-level predictions obtained during outer-fold cross-validation are presented for early-fusion CNN1D classifiers trained on feature sets derived from all combinations of strains displaying prognostic capabilities (A0, A1, A5, A15 and A19). For each strain, the feature set used was the one associated with the highest balanced accuracy in the corresponding individual-strain classifier: (i) time series and first derivative (both not normalized) combined with growth-parameter conditioning through feature-wise linear modulation (FiLM) for A0, (ii) time series combined with the first derivative (both normalized) for A1, (iii) time series only (not normalized) for A5, (iv) time series combined with the first derivative (both not normalized) for A15 and (v) time series combined with the first derivative (both normalized) for A19. Green dots indicate correct predictions, yellow dots indicate mild patients misclassified as severe, and orange dots indicate severe patients misclassified as mild.

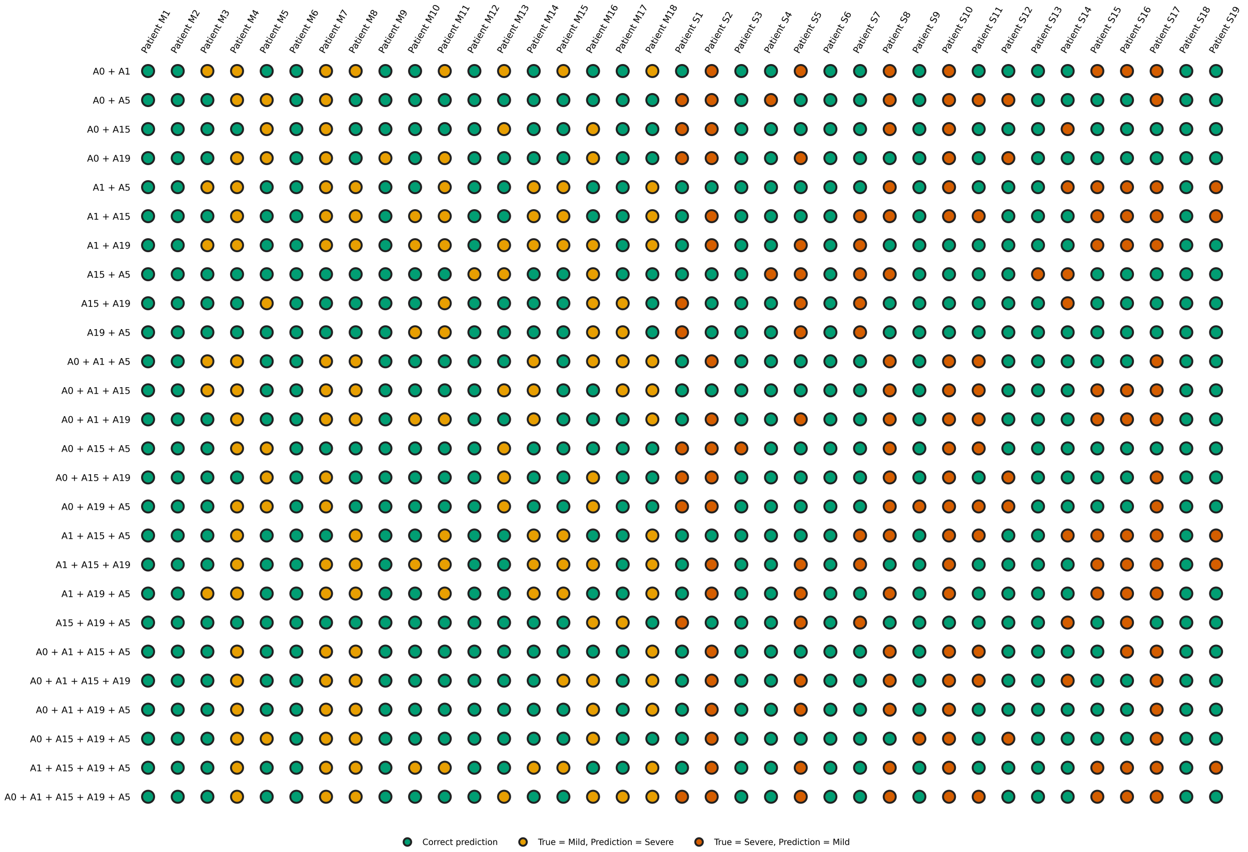


Supplementary Figure 10: Outer-fold cross-validation patient-level predictions of early-fusion temporal convolutional network (TCN) models for prognostic classification.

Patient-level predictions obtained during outer-fold cross-validation are presented for early-fusion TCN classifiers trained on feature sets derived from all combinations of strains displaying prognostic capabilities (A0, A1, A5, A15 and A19). For each strain, the feature set used was the one associated with the highest balanced accuracy in the corresponding individual-strain classifier: (i) time series and first derivative (both not normalized) combined with growth-parameter conditioning through feature-wise linear modulation (FiLM) for A0, (ii) time series combined with the first derivative (both normalized) for A1, (iii) time series only (not normalized) for A5, (iv) time series combined with the first derivative (both not normalized) for A15 and (v) time series combined with the first derivative (both normalized) for A19. Green dots indicate correct predictions, yellow dots indicate mild patients misclassified as severe, and orange dots indicate severe patients misclassified as mild.

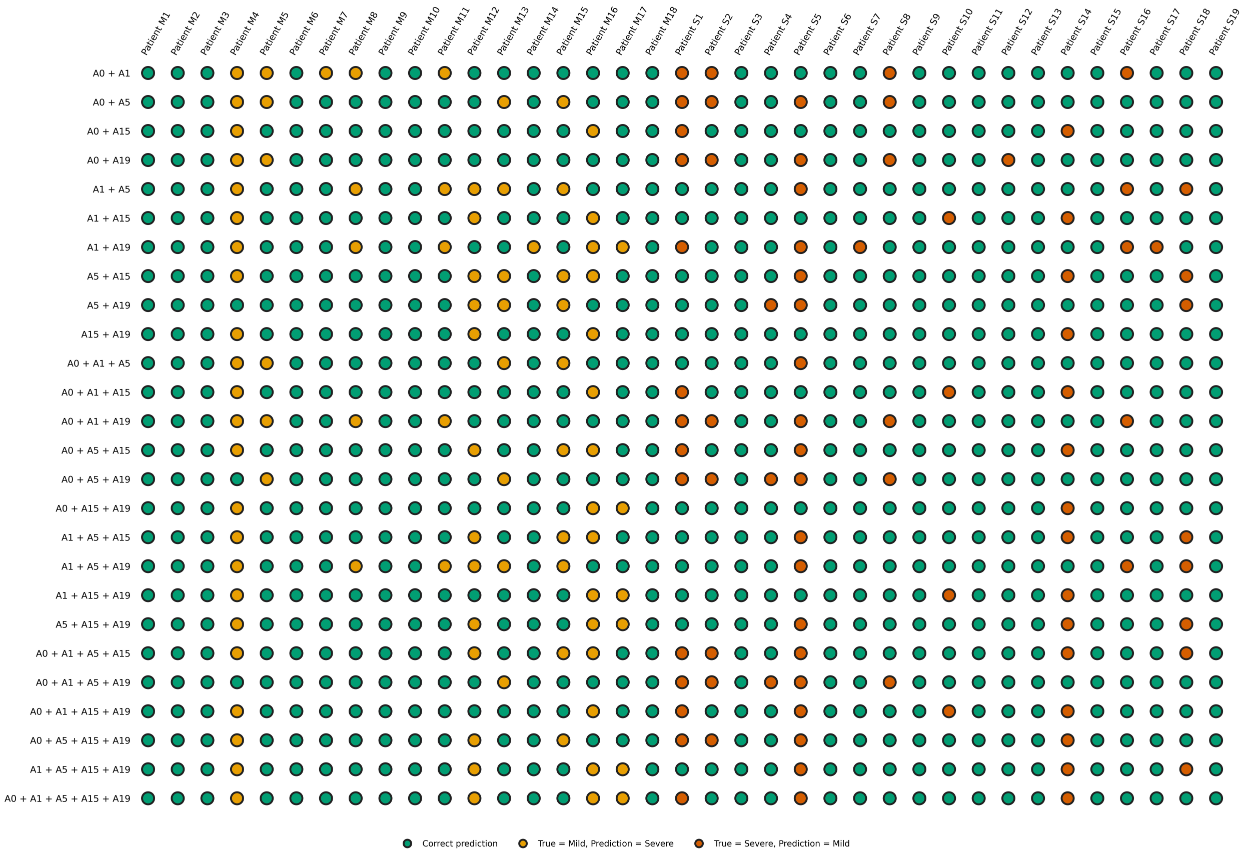

Supplementary Figure 11: Outer-fold cross-validation patient-level predictions of weighted soft-voting late-fusion models for prognostic classification.

Patient-level predictions obtained during outer-fold cross-validation are presented for weighted soft-voting late-fusion ensemble built from individual-strain models. Individual-strain models are trained on the feature set yielding the highest balanced accuracy for the corresponding strain: (i) time series and first derivative (both not normalized) combined with growth-parameter conditioning through feature-wise linear modulation (FiLM) for A0, (ii) time series combined with the first derivative (both normalized) for A1, (iii) time series only (not normalized) for A5, (iv) time series combined with the first derivative (both not normalized) for A15 and (v) time series combined with the first derivative (both normalized) for A19. Green dots indicate correct predictions, yellow dots indicate mild patients misclassified as severe, and orange dots indicate severe patients misclassified as mild.

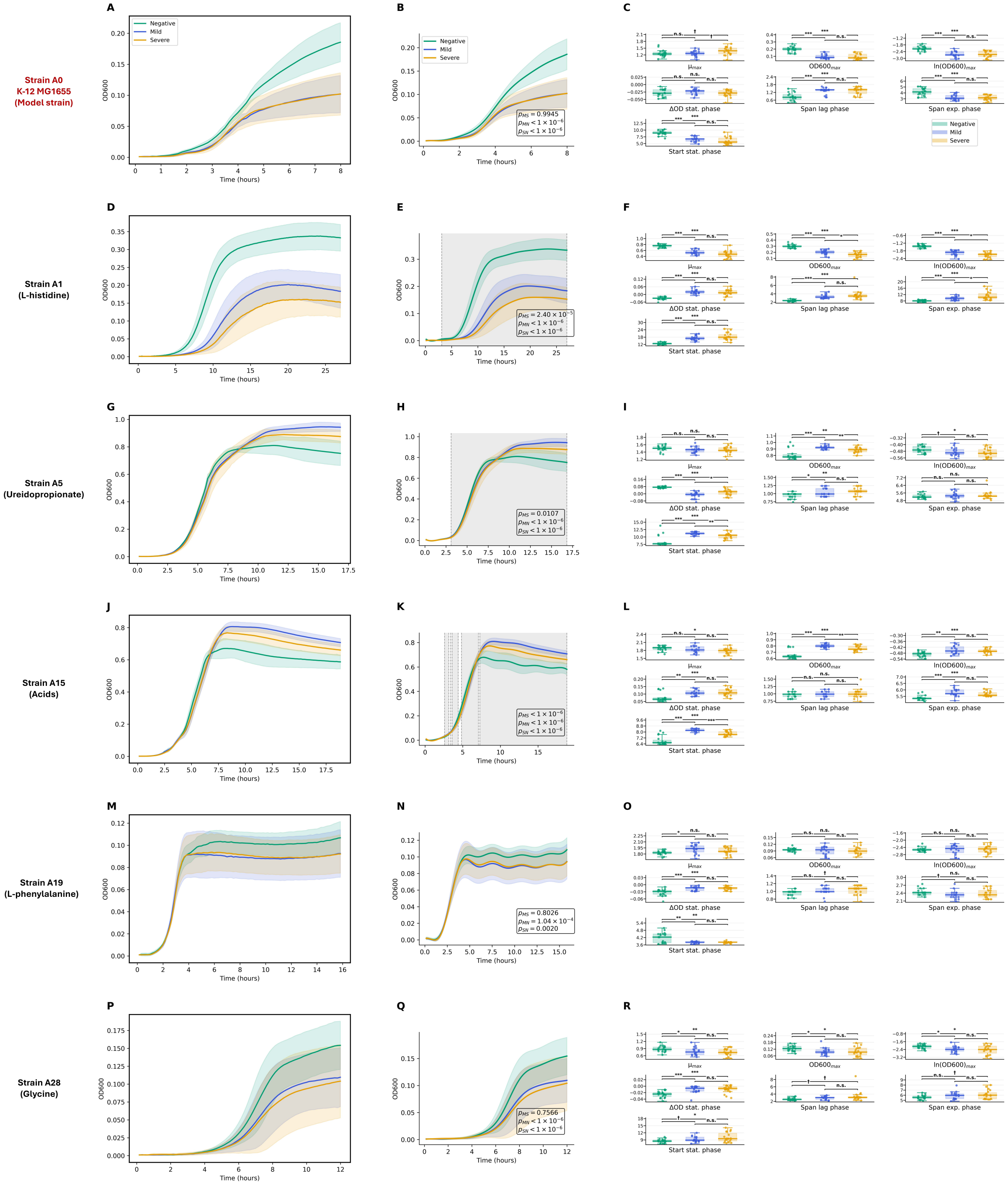


Supplementary Figure 12: Confirmation of group-wise mean trajectory differences between mild, severe and negative classes for the five selected metabolic sensors.

For each sensor candidate, mean growth curves for the mild ($n = 18$), severe ($n = 19$) and negative ($n = 19$) patient samples are illustrated in the left section of the first column of panels (A, D, G, J, M, P). Mild curves are colored in blue, severe curves in orange, negative curves in green and positive curves in red. Shaded areas represent standard deviations. The second column of panels (B, E, H, K, N, Q) depicts GAMM-fitted mild, severe and negative growth curves, together with corresponding $p$-values for all pairwise group comparisons ($\alpha = 0.1$). $p$-values were corrected using Benjamini-Hochberg procedure [2] to account for multiple testing. Shaded areas represent standard deviations, and hatched areas indicate 95% confidence intervals (not readily visible in most plots owing to their narrow range). Grey windows in the GAMM representations indicate time intervals showing significant differences between conditions. The last column of panels (C, F, I, L, O, R) shows boxplots for each growth parameter. Each dot represents a growth curve acquired on an individual patient sample. $p$-values (corrected using Benjamini-Hochberg procedure [2] to account for multiple testing) for all pairwise group comparisons (Welch's $t$-test, $\alpha = 0.1$) are indicated as follows: $p < 0.001$ (***), $p < 0.01$ (**), $p < 0.05$ (*), $p < 0.1$ (†) and $p > 0.1$ or non-significant ($n.s.$).

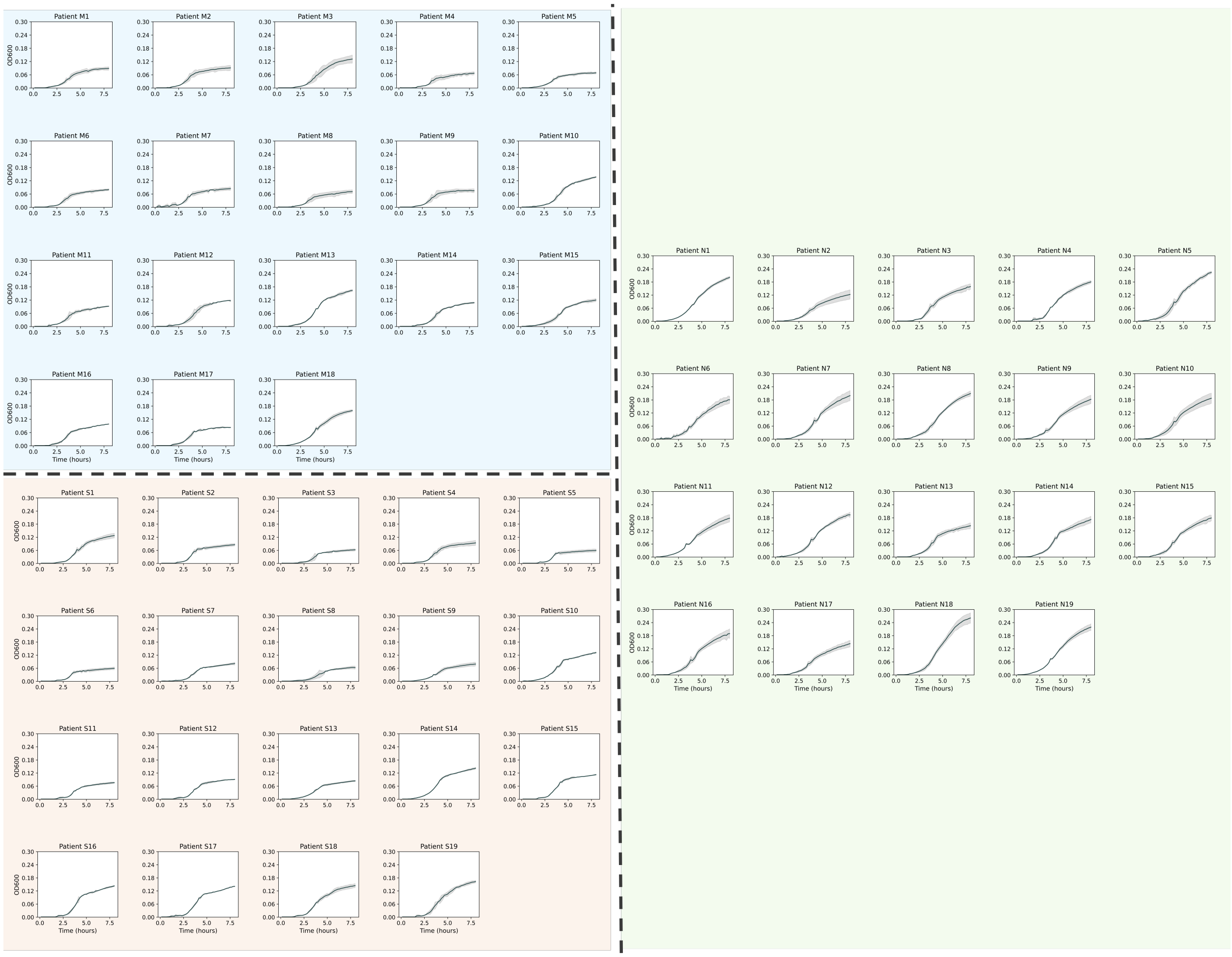

Supplementary Figure 13: Growth curves of A0 strain cultured in plasma samples from the validation cohort. Standard deviations are indicated by shaded areas. Growth curves in the upper-left (blue), lower-left (orange), and right (green) sections correspond to mild ($n = 18$), severe ($n = 19$), and negative subjects ($n = 19$), respectively.

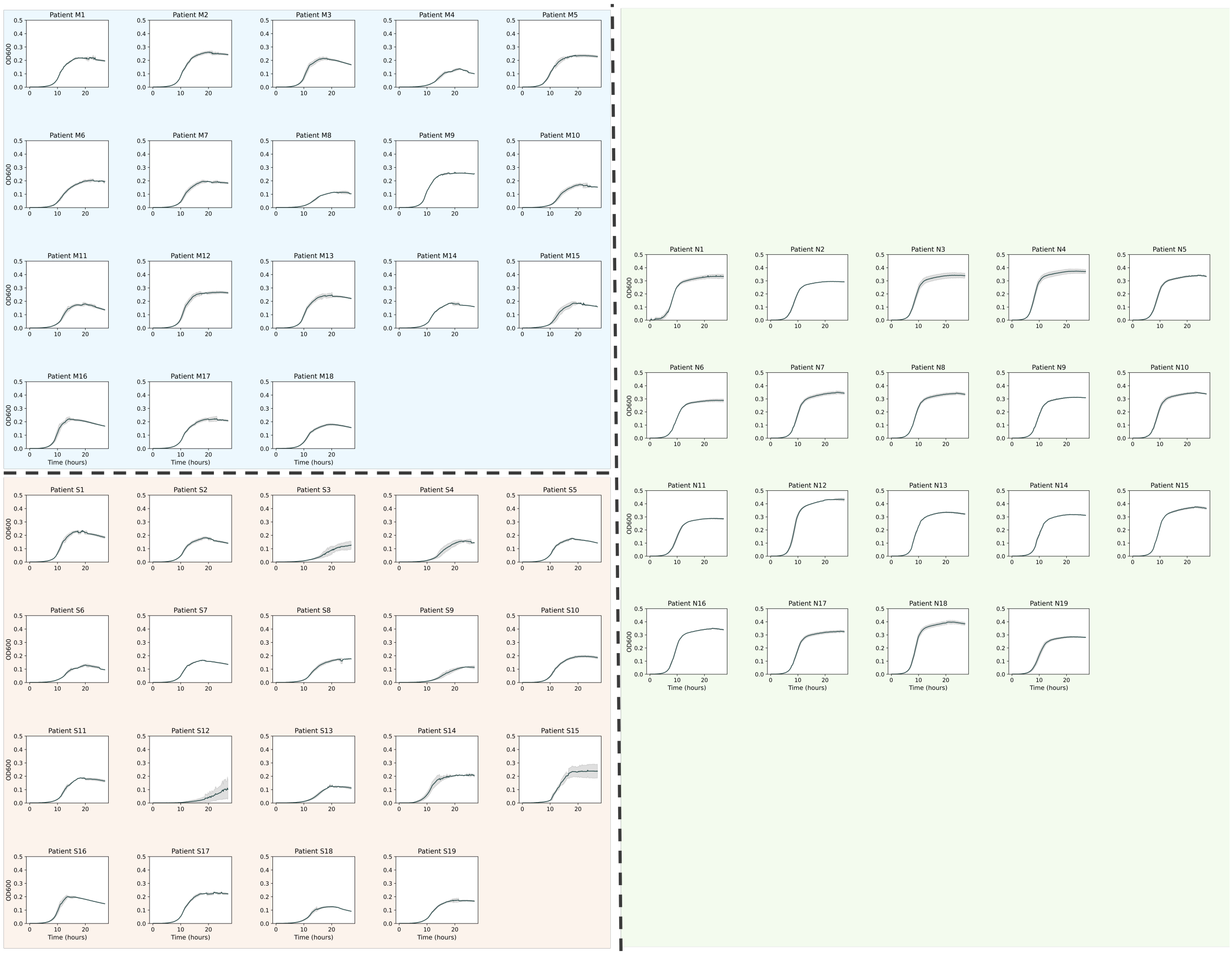


Supplementary Figure 14: Growth curves of A1 strain cultured in plasma samples from the validation cohort. Standard deviations are indicated by shaded areas. Growth curves in the upper-left (blue), lower-left (orange), and right (green) sections correspond to mild ($n = 18$), severe ($n = 19$), and negative subjects ($n = 19$), respectively.

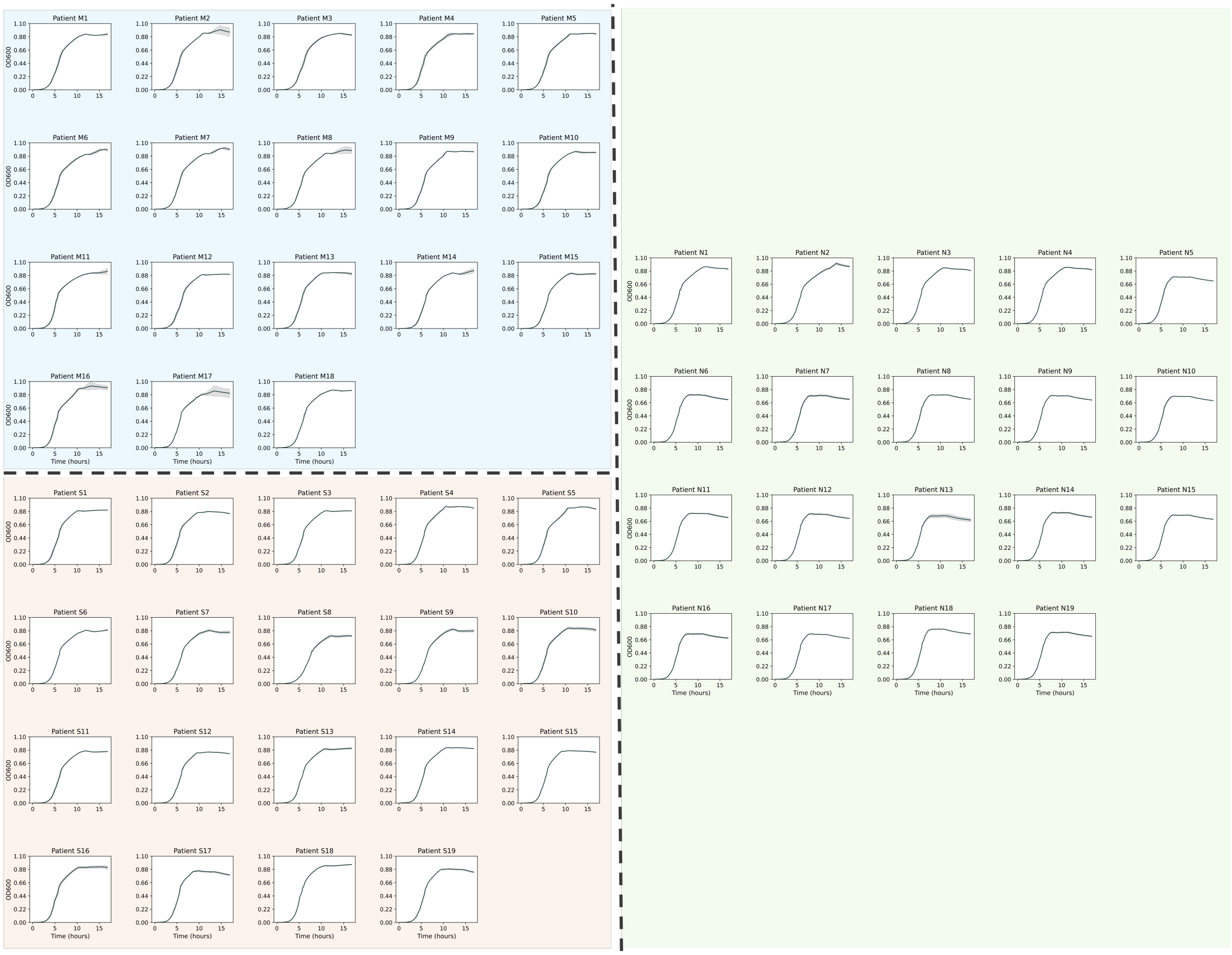


Supplementary Figure 15: Growth curves of A5 strain cultured in plasma samples from the validation cohort. Standard deviations are indicated by shaded areas. Growth curves in the upper-left (blue), lower-left (orange), and right (green) sections correspond to mild ($n = 18$), severe ($n = 19$), and negative subjects ($n = 19$), respectively.

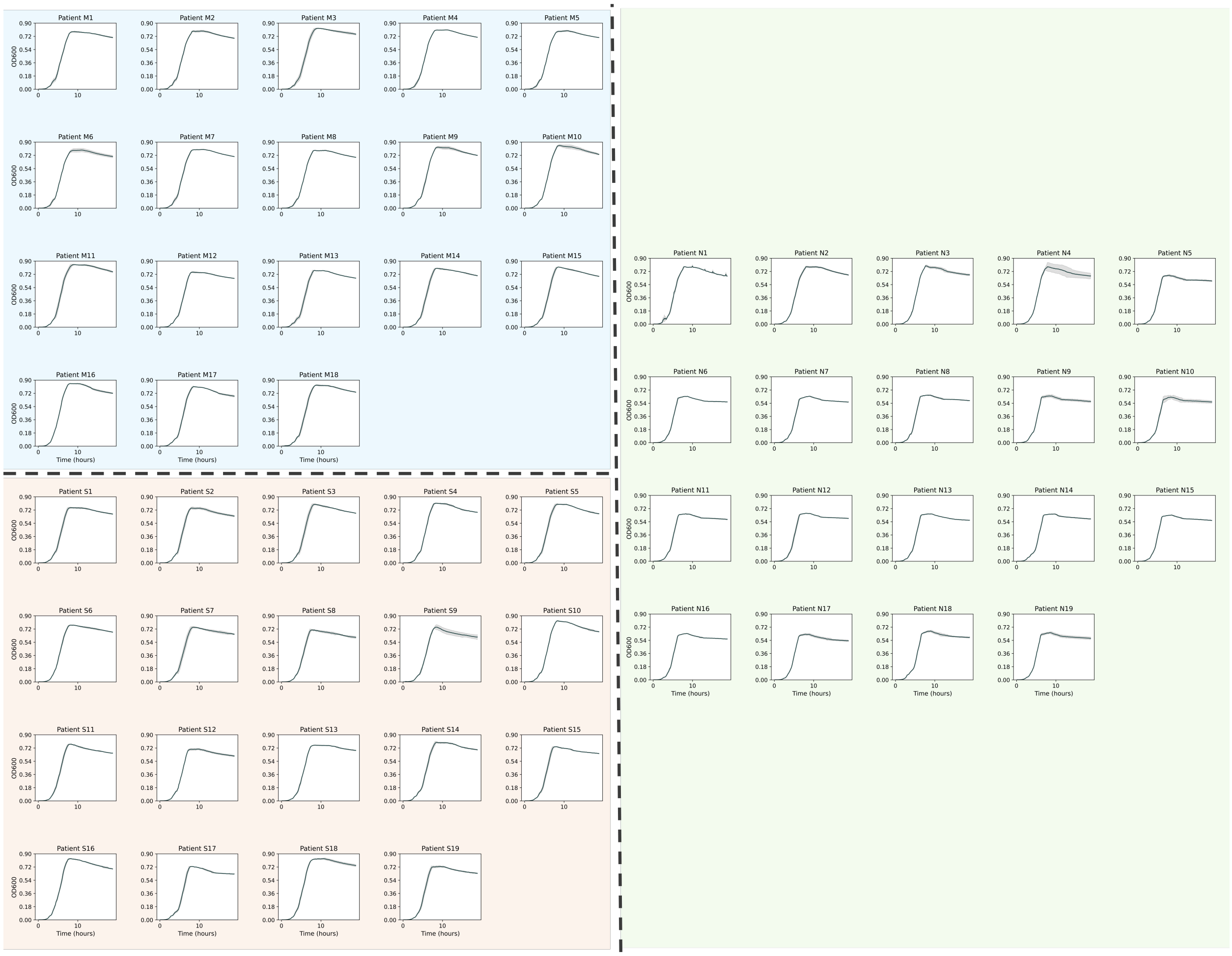


Supplementary Figure 16: Growth curves of A15 strain cultured in plasma samples from the validation cohort.

Standard deviations are indicated by shaded areas. Growth curves in the upper-left (blue), lower-left (orange), and right (green) sections correspond to mild ($n = 18$), severe ($n = 19$), and negative subjects ($n = 19$), respectively.

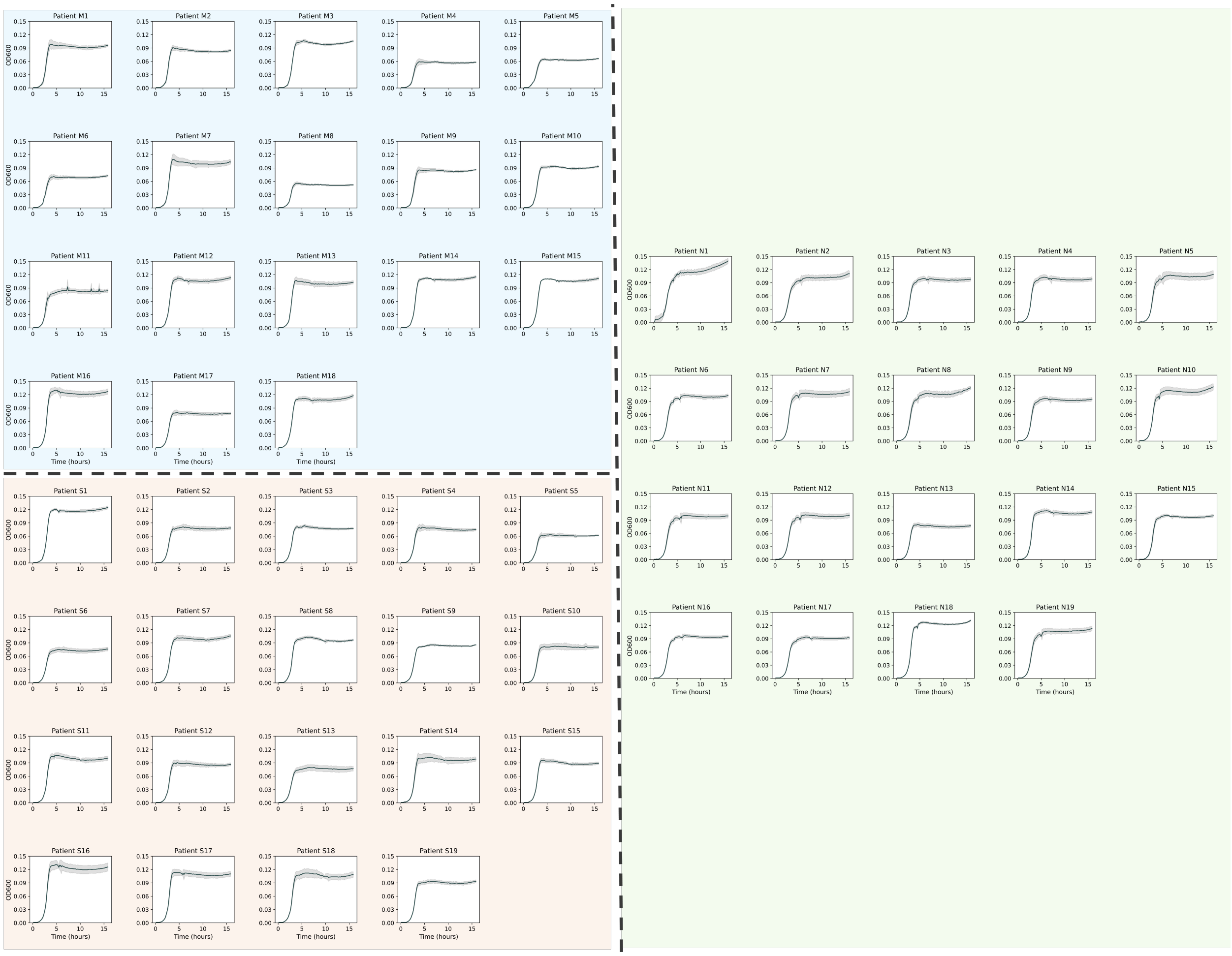


Supplementary Figure 17: Growth curves of A19 strain cultured in plasma samples from the validation cohort.

Standard deviations are indicated by shaded areas. Growth curves in the upper-left (blue), lower-left (orange), and right (green) sections correspond to mild ($n = 18$), severe ($n = 19$), and negative subjects ($n = 19$), respectively.

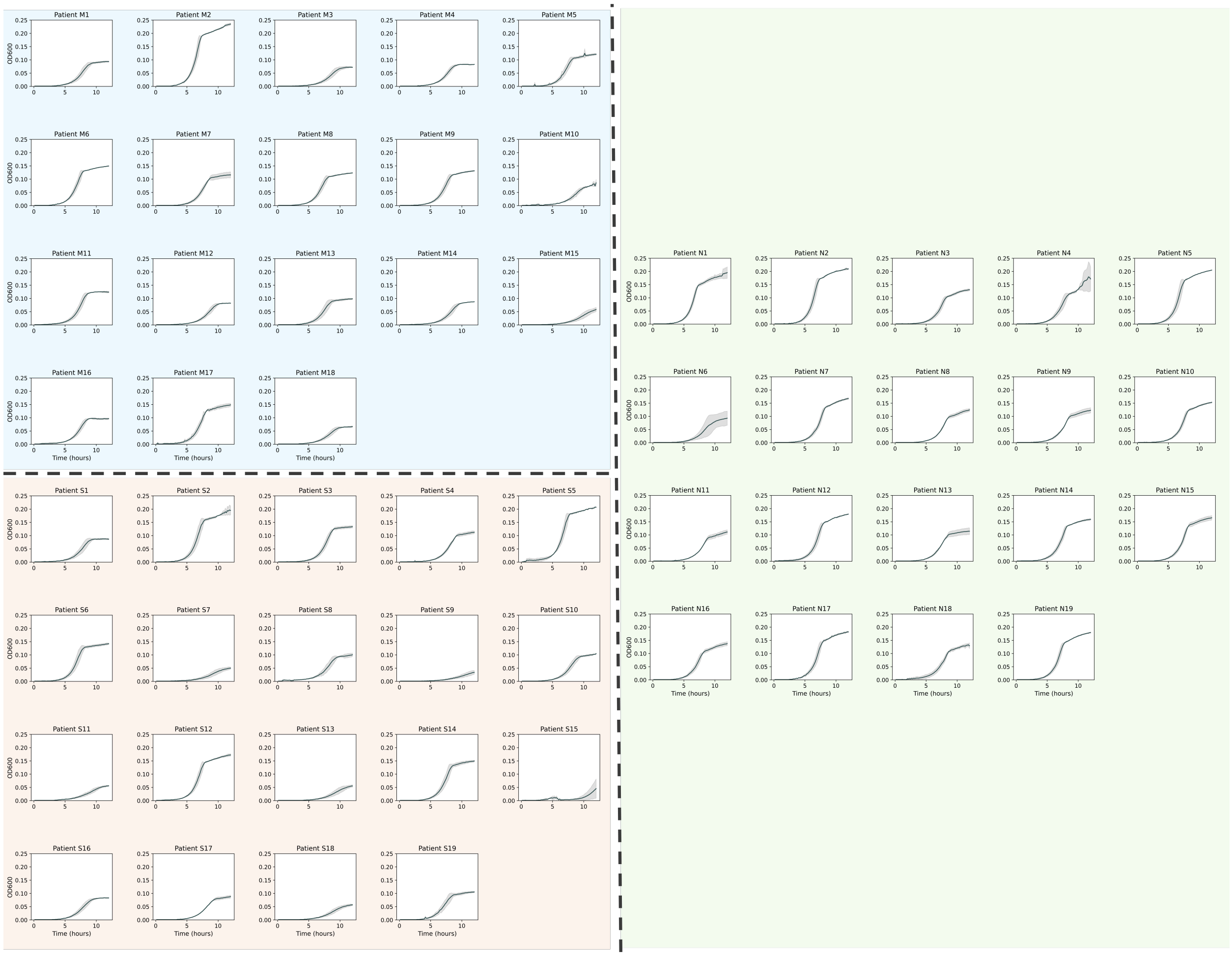

Supplementary Figure 18: Growth curves of A28 strain cultured in plasma samples from the validation cohort. Standard deviations are indicated by shaded areas. Growth curves in the upper-left (blue), lower-left (orange), and right (green) sections correspond to mild ($n = 18$), severe ($n = 19$), and negative subjects ($n = 19$), respectively.

### References


(1) Lin, X.; Zhang, D. *Journal of the Royal Statistical Society Series B: Statistical Methodology* **1999**, *61*, 381–400.

(2) Benjamini, Y.; Hochberg, Y. *Journal of the Royal Statistical Society Series B: Statistical Methodology* **1995**, *57*, 289–300.

(3) Perez, E.; Strub, F.; de Vries, H.; Dumoulin, V.; Courville, A. FiLM: Visual Reasoning with a General Conditioning Layer, Version Number: 2, 2017.
