## Supplementary material for "Engineering growth-coupled metabolic biosensors for disease prognosis and diagnosis using full growth trajectories": README.txt

README

This folder contains the first part of the supplementary data associated with Ahavi et al. It compiles the results of the sensor screening phase and includes:

(i) a file named "Sensor screening statistics". It contains the measured OD600 values corresponding to the uninterpretable growth curves and the OD600 values acquired over time as well as the growth parameters of each screened strain grown on the three biological replicates (plasma sample pools).

(ii) a folder named "GAMM_model selection" containing all results of the GAMM (Generalized Additive Mixed Model) model selection procedure. "MS" stands for "Mild vs severe", "NP" for "Negative vs positive", "MN" for "Mild vs Negative", "NS" for "Negative vs severe" and "three_groups" for "Mild vs severe vs negative".

(iii) a folder named "GAMM_sensor screening" containing all results of the statistical tests for all classification problems based on the selected GAMMs. The same problem abbreviations are used.
