## Supplementary material for "Engineering growth-coupled metabolic biosensors for disease prognosis and diagnosis using full growth trajectories": README.txt

README

This folder contains the second part of the supplementary data associated with Ahavi et al. It compiles the results of the sensor validation phase and includes:

(i) a folder named "Growth curves". Each file summarizes the OD600 values measured over time for each strain grown on each patient sample from the validation cohort. The first sheet of each file reports both technical replicates for each patient, while the second sheet comprises the mean and standard deviation at each time point.

(ii) a folder named "Growth parameters". Each file summarizes the seven growth parameters extracted from the growth curves of each patient (µmax, OD600max, ln(OD600)max, span of the lag phase, span of the exponential phase, start of the stationary phase, OD600max - OD600final difference). Each file contains four sheets: "Replicate 1" and "Replicate 2" for both technical replicates per patient and then "Mean" and "Standard deviation".

(iii) a file named "Welch_t-test". It contains the results of Welch's t-tests conducted on the growth parameters for all three classification problems (Mild vs severe, Negative vs positive, Mild vs severe vs negative).

(iv) a folder named "GAMM_model selection" containing all results of the GAMM (Generalized Additive Mixed Model) model selection procedure. "MS" stands for "Mild vs severe", "NP" for "Negative vs positive", "MN" for "Mild vs Negative", "NS" for "Negative vs severe" and "three_groups" for "Mild vs severe vs negative".

(v) a folder named "GAMM_sensor validation" containing all results of the statistical tests based on the selected GAMMs. The same problem abbreviations are used.
