## Supplementary material for "Engineering growth-coupled metabolic biosensors for disease prognosis and diagnosis using full growth trajectories": README.txt

README

This folder contains nine excel files reporting classification performance metrics, raw folds and patient predictions of different models developed in this study:

(i) "Summary - Best model" report results corresponding to the model yielding the highest accuracy for each strain and each classification problem (prognosis, diagnosis, multiclass) across the four input feature sets considered.

(ii) "Summary - Seven growth parameters - With feature selection", "Summary - Seven growth parameters - Without feature selection" and "Summary - Two growth parameters" report results of the 4 classifiers tested (SVM, LogReg, XGBoost, VotingEnsemble) for growth-parameter-based classification across all strains, using whether two or seven growth parameters as input set and applying or not feature selection.

(iii) "Summary - Time series", "Summary - Time series + derivatives", "Summary - Time series (best input set) + growth parameters (FiLM)" recapitulates results corresponding to the model associated with the highest balanced accuracy (CNN1D or TCN) for time-series classification across all strains and input sets considered.

(iv) "Summary - Multistrain models" report results of all early- and late-fusion models.
